# Mitochondrial ATP Synthase Subunit d, a Component of the Peripheral Stalk, is Essential for Growth and Heat Stress Tolerance in *Arabidopsis thaliana*

**DOI:** 10.1101/2021.02.02.429459

**Authors:** Tianxiang Liu, Jesse Arsenault, Elizabeth Vierling, Minsoo Kim

**Author notes:** Corresponding author: Minsoo Kim.

## Abstract

As rapid changes in climate threaten global crop yields, an understanding of plant heat stress tolerance is increasingly relevant. Heat stress tolerance involves the coordinated action of many cellular processes and is particularly energy demanding. We acquired a knockout mutant and generated knockdown lines in *Arabidopsis thaliana* of the d subunit of mitochondrial ATP synthase (gene name: *ATPQ*, AT3G52300, referred to hereafter as *ATPd*), a subunit of the peripheral stalk, and used these to investigate the phenotypic significance of this subunit in normal growth and heat stress tolerance. Homozygous knockout mutants for *ATPd* could not be obtained due to gametophytic defects, while heterozygotes possess no visible phenotype. Therefore, we used RNAi to create knockdown plant lines for further studies. Proteomic analysis and blue native gels revealed that *ATPd* downregulation impairs only subunits of the mitochondrial ATP synthase (complex V of the electron transport chain). Knockdown plants were more sensitive to heat stress, had abnormal leaf morphology, and were severely slow growing compared to wild type. These results indicate that *ATPd* plays a crucial role in proper function of the mitochondrial ATP synthase holoenzyme, which, when reduced, leads to wide-ranging defects in energy-demanding cellular processes. In knockdown plants, more hydrogen peroxide accumulated and mitochondrial dysfunction stimulon (MDS) genes were activated. These data establish the essential structural role of *ATPd* and provide new information about complex V assembly and quality control, as well as support the importance of mitochondrial respiration in normal plant growth and heat stress tolerance.

**SIGNIFICANCE STATEMENT:** The energy converter, mitochondrial ATP synthase, is critical for all organisms, but the functional importance of ATP synthase subunit d remains largely unknown in plants. We demonstrate the contributions of subunit d to plant growth, development, and heat stress tolerance, as well as to ATP synthase stability, ROS signaling and mitochondrial dysfunction regulation.

## INTRODUCTION

The mitochondrial inner membrane-located ATP synthase (or F_1_F_O_-ATP synthase) catalyzes the terminal step of oxidative phosphorylation (OXPHOS), producing energy-rich ATP from the electron transport-generated proton gradient. The main subcomplexes of ATP synthase are the soluble F_1_ and membrane-embedded F_O_, which perform the important functions of catalysis and proton-pumping, respectively, along with the central stalk and the peripheral stalk (PS) that connect F_1_ and F_O_ (Welch *et al.*, 2011; Artika, 2019; Mühleip *et al.*, 2019; Kühlbrandt, 2019; Pinke *et al.*, 2020). The basic subunit composition and structure of mitochondrial ATP synthase are conserved in eukaryotes and have many similarities to the bacterial counterpart (Zancani *et al.*, 2020). The F_1_ sector consists of the catalytic head composed of three αβ dimers and the central stalk containing subunits γ, δ and ε. The composition of F_O_ is slightly different in plants compared to yeast and mammals. In *Arabidopsis thaliana*, the F_O_ sector consists of the motor with subunits a, c, e, f, g, and 8 (ATP8 or ORFB), and the PS comprises subunits b, d and OSCP. Notably, the PS in plants lack subunit h (F_6_) that is present in yeast and animals. The *A. thaliana* F_O_ additionally has plant-specific F_A_d and 6 kDa subunits (Li *et al.*, 2010; Brugière *et al.*, 2004). While only two subunits (a and 8) are encoded in the human mitochondrial genome, three additional subunits (α, b, c) are encoded in *A. thaliana* mitochondrial DNA and synthesized in the mitochondrial matrix. Mitochondrial ATP synthases form dimers, which were first purified from yeast (Arnold *et al.*, 1998). Remarkably, these dimers oligomerize into rows inducing curvature of the inner mitochondrial membrane critical for the formation of mitochondrial cristae (Paumard *et al.*, 2002; Dudkina *et al.*, 2010; Blum *et al.*, 2019).

Energy is the basis for organisms to perform all life activities. Therefore, abnormalities in ATP synthase subunits cause severe growth defects and usually lethality in *A. thaliana* (Robison *et al.*, 2009; Li *et al.*, 2010; Geisler *et al.*, 2012). In many crop species, aberrations in all five mitochondrial genes (*ATP1* (subunit α), *ATP4* (subunit b), *ATP6* (subunit a), *ATP8* (subunit 8) and *ATP9* (subunit c)) are associated with cytoplasmic male sterility (CMS), a trait where the functional male reproductive structures fail to develop correctly, resulting in the partial or total loss of pollen development (reviewed in Horn *et al.*, 2014; Chen and Liu, 2014). Mutations in nuclear genes encoding mitochondrial ATP synthase subunits also result in male sterility. Knocking out nuclear genes encoding central stalk subunit δ and plant specific F_A_d cause male sterility as well as female defects in *A. thaliana* (Li *et al.*, 2010; Geisler *et al.*, 2012). T-DNA insertional mutants of subunit OSCP were screened but no homozygous mutants were obtained due to gametophytic lethality (Moore *et al.*, 2003). Therefore, many ATP synthase subunits are essential for plant growth and development.

In addition to the critical role of mitochondria as ATP generators, mitochondria play key roles in the reception of stress signals and responses (Rasmusson and Møller, 2011; Jacoby *et al.*, 2012). Under conditions that impair mitochondrial electron transport, such as oxidative and heat stress, plant mitochondria produce reactive oxygen species (ROS) (Møller, 2001; Wang *et al.*, 2018). ROS can act as a signal to trigger mitochondrial retrograde regulation (MRR) activating mitochondrial dysfunction stimulon (MDS) genes (De Clercq *et al.*, 2013; Ng *et al.*, 2013; Shapiguzov *et al.*, 2019). Products of MDS genes, such as AOX1a, OM66, MGE1, and ANAC013, likely help repair dysfunctional mitochondria. Transient inhibition of complex V with oligomycin treatment also induces MDS genes such as AOX1a and ANAC013 (Clifton *et al.*, 2005; Schwarzländer *et al.*, 2012). Notably, hierarchical clustering of RNA expression profiles put oligomycin treatment in the same group with rotenone (complex I inhibitor) and antimycin A (complex III inhibitor) treatments, reflecting a common response to OXPHOS inhibition (Schwarzländer *et al.*, 2012). However, it is not known to what extent chronic defects of complex V trigger MRR in plants.

Subunit d of the PS of the mitochondrial ATP synthase has been investigated in humans and fruit flies, but remains largely uncharacterized in plants. One plant proteomics study indicated that ATPd decreased in response to hydrogen peroxide treatment (Tan *et al.*, 2012). In other organisms, downregulation of this subunit greatly impairs ATP synthase function and results in major phenotypic alterations in processes related to metabolic efficiency. RNAi knockdowns of ATPd in Drosophila significantly increased oxidative stress tolerance, decreased protein aggregation, and improved lifespan in fruit flies fed with a low carbohydrate-to-protein diet (Sun *et al.*, 2014). In humans, knockdowns of ATPd inhibited the assembly of the holoenzyme (Fujikawa *et al.*, 2015; He *et al.*, 2020). Because ATP synthase is central to energy homeostasis, additional studies of ATPd in plants may yield further insights into the involvement of mitochondrial ATP synthase in growth and stress tolerance. We investigated the biological function of mitochondrial ATP synthase subunit d in *A. thaliana*. We show that knockout of ATPd leads to male sterility as well as female defects. Knockdown of ATPd by RNAi exerts pleiotropic effects on plant growth and development including severe growth retardation, late flowering, floral defects, and reduced fertility. Label-free quantitative proteomics analysis of mitochondria from ATPd knockdown plants suggests decreased stability of complex V assembly intermediates but has surprisingly little effect on other OXPHOS components. Furthermore, enhanced heat sensitivity of *ATPd* RNAi plants is correlated with increased ROS accumulation and expression of a number of MDS genes.

## RESULTS

### Absence mitochondrial ATP synthase subunitd causes male sterility and female defects

The *ATPQ* gene (AT3G52300, *ATPd*) encoding mitochondrial ATP synthase subunit d contains 5 exons (Figure 1a) specifying a protein of 168 amino acids. Alignment of ATPd amino acid sequences from *A. thaliana* and rice shows extensive conservation (76% identity), but limited conservation with human (27% identity) and yeast (24% identity) homologues (Figure S1), suggesting minimal specific sequence requirements for its function in the PS. To investigate the biological role of ATPd on plant growth and development, we obtained T-DNA insertion mutants for the *ATPd* gene from the SAIL mutant collection (Sessions *et al.*, 2002). Two lines were identified, of which one contained a T-DNA in the second intron (SAIL_97_F05, *atpd-1*), and the other had an insert in the fourth intron (SAIL_515_E09, *atpd-2*; Figure 1a). We were able to identify homozygous *atpd-2* plants (Figure 1b) without any decrease in transcript level (Figure 1d) implying that the T-DNA insert in the fourth intron was properly spliced out. By contrast, no homozygous *atpd-1* plants were obtained indicating that the T-DNA in the second intron disrupted gene expression. Heterozygous *atpd-1*/+ plants did not show any obvious growth defects compared to wild-type siblings (Figure 1c). To confirm that the T-DNA in *atpd-1* causes lethality, a genomic DNA construct under the constitutive 35S promoter was introduced into *atpd-1*/+ plants. Both heterozygous and homozygous *atpd-1* plants were obtained in the T2 generation and no significant growth differences were observed between complemented *atpd-1* and wild type (Col-0) (Figure 1e and f), confirming that the lack of ATPd causes lethality.

**Figure 1.**
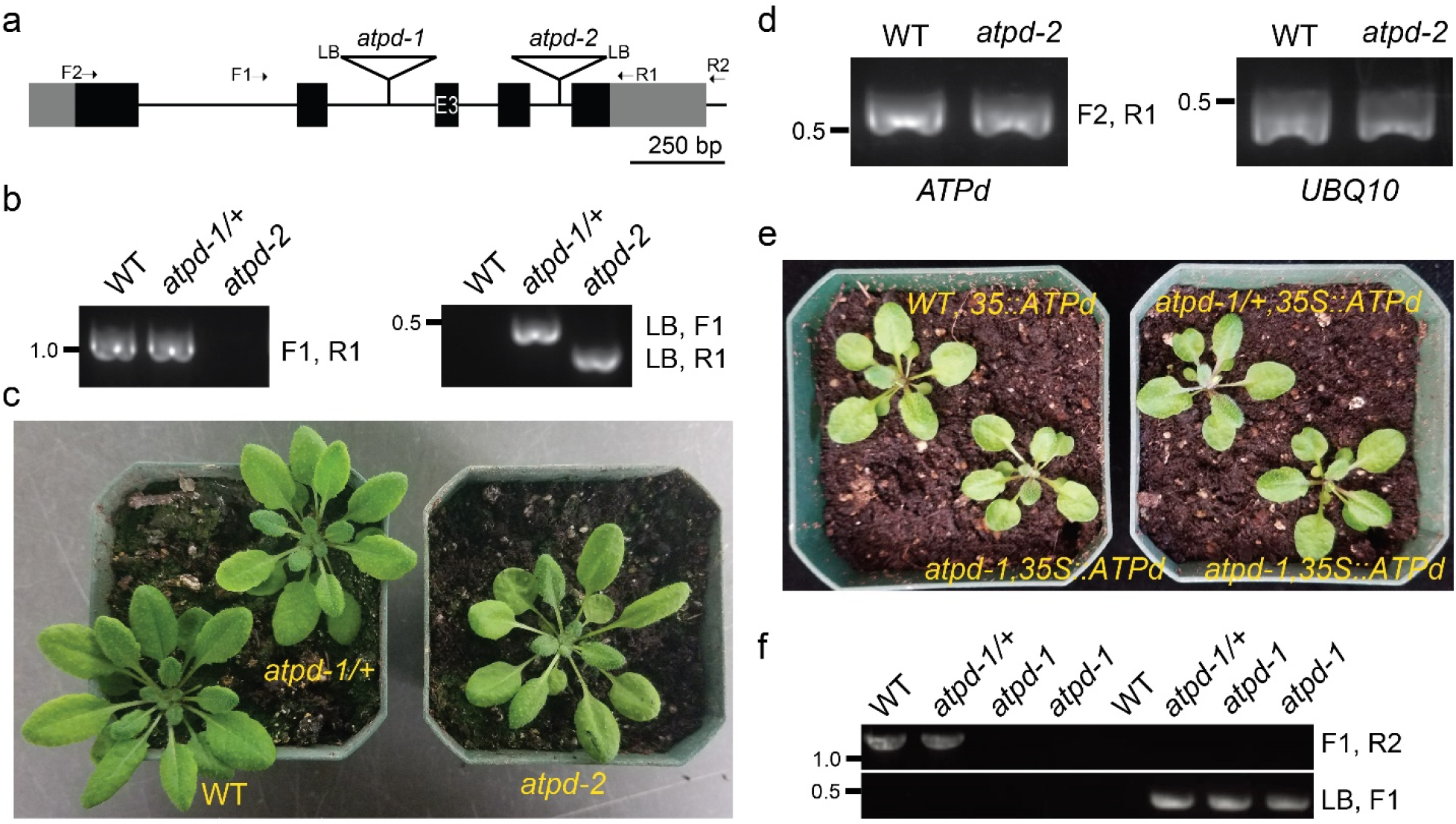
ATP synthase subunit d is essential for plant growth. (a) Gene structure of *ATPd* and positions of T-DNA insertions in *atpd-1* (SAIL_97_F05) and *atpd-2* (SAIL_515_E09) mutants. F1, F2, LB, R1 and R2 indicate primers used in genotyping and RT-PCR. Black and grey boxes indicate the exons and UTRs, respectively. (b) PCR genotyping results show that homozygous *atpd-1* mutants cannot be obtained from *atpd-1*/+ plants. Homozygous *atpd-2* mutant plants can be obtained. (c) Plant growth phenotypes of *aptd-1*/+ and *atpd-2*. (d) RT-PCR analysis show normal expression of *ATPd* in the *atpd-2* mutant. (e) Genomic DNA of *ATPd* under 35S promoter complements *atpd-1*. (f) PCR genotyping analysis shows that homozygous *atpd-1* mutants can be obtained from *35S::ATPd* T2 plants. The R2 primer anneals further downstream of the 3’ end of the construct so that only the endogenous *ATPd* gene can be amplified.

It has been reported that mutations in subunits of the ATP synthase complex cause male sterility and female defects (Li *et al.*, 2010; Geisler *et al.*, 2012). Therefore, we investigated the segregation ratio of progenies of *atpd-1*/+ plants. A segregation ratio of 1:2:1 of each genotype AA, Aa, and aa would be expected if there were no defect caused by the mutation. Out of 95 progenies genotyped, we observed a ratio of approximately 5:1:0 (Figure 2a), indicating that the T-DNA allele was not fully transmitted. To determine transmission efficiencies of the T-DNA insertion, reciprocal backcrosses between heterozygous *atpd-1*/+ and wild-type plants were performed (Figure 2a). If the haploid gametophytes were unaffected by the T-DNA insertion, a 1:1 genotypic ratio of wild type to heterozygous progeny would be expected. Fertilization of the wild-type female with the *atpd-1*/+ pollen produced only wild-type plants, indicating that the mutation could not be transmitted through mutant pollen. The reciprocal cross between the *atpd-1*/+ female and the wild-type produced only 18% *atpd-1*/+ plants, indicating that the lack of *ATPd* gene also causes defects in the female. These results are similar to observations made in knockout mutants in subunits δ and F_A_d of the ATP synthase (Li *et al.*, 2010, Geisler *et al.*, 2012).

**Figure 2.**
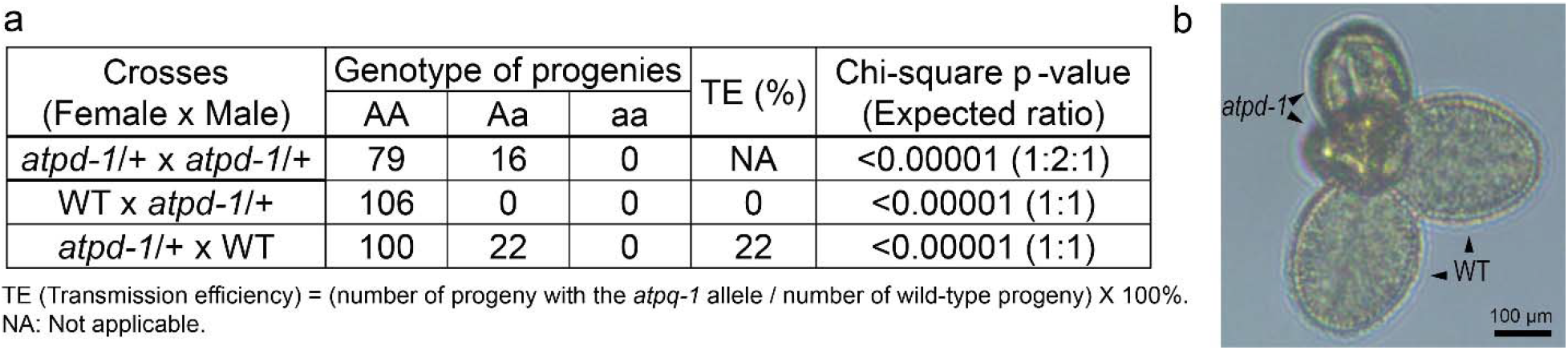
Lack of ATP synthase subunit d causes male sterility and a female defect. (a) Genetic analysis of F1 progenies of *atpd-1*/+ self-crosses and reciprocal backcrosses with the wild type. Chi-square test was performed to compare the observed ratios with the expected ratios in parenthesis. (b) Light microscopy image of a mature pollen tetrad from *atpd-1/+ qrt2* mutant plants.

To further explore the effects of *atpd-1* on male gametophyte development, *atpd-1*/+ plants were crossed into the *quartet 2* (*qrt2*) mutant background (SALK_026522C). *QUARTET2 (QRT2)* encodes a polygalacturonase in *A. thaliana* that, when knocked out, inhibits the separation of developing pollen grain tetrads (Ogawa *et al.*, 2009). These tetrahedral arrangements of haploid cells can be visualized, and because they descend from the same mother cell, will each possess either the wild type or *atpd-1* allele in a 1:1 ratio. If a mutant allele negatively impacts development, there will often be visible defects in the affected cells. Seeds from *atpd-1*/+ *qrt2* background were grown on soil and genotyped to identify plants with the *atpd-1* T-DNA insertion. Pollen was isolated from the anthers of open flowers and visualized under a light microscope. Pollen grains clearly exhibited major deformities in two of the tetrad cells (Figure 2b), indicating that the mutation causes defects in pollen development.

### Growth is severely reduced with down-regulation of ATPd

To assess the effects of reduced expression of ATPd, we generated RNAi knockdown lines by transforming wild type *A. thaliana* plants with a 300 bp fragment from near the 3’-end of *ATPd* cDNA in sense and antisense direction under the 35S promoter. Five independent knockdown lines were selected for initial phenotypic characterization. All five knockdown lines (RNAi lines 1 to 5) displayed severe growth retardation under long day growth conditions (16h light / 8h dark) (Figure 3a and S2a), typical of plants with mitochondrial dysfunction, while the empty vector control (EV) plants grew like the wild-type Columbia-0. In addition, RNAi lines had malformed leaves and flowers, featuring irregular or “crinkled” edges and abnormally small leaf sizes compared to those of the wild type (Figure 3a, S2a and b). They also have short roots and fewer root hairs, short stems, short siliques and misshapen flowers and floral organs resulting in severe fertility defects (Figure 3a-e). Among the five RNAi lines, RNAi-2 had a slightly milder phenotype (Figure 3a-e).

**Figure 3.**
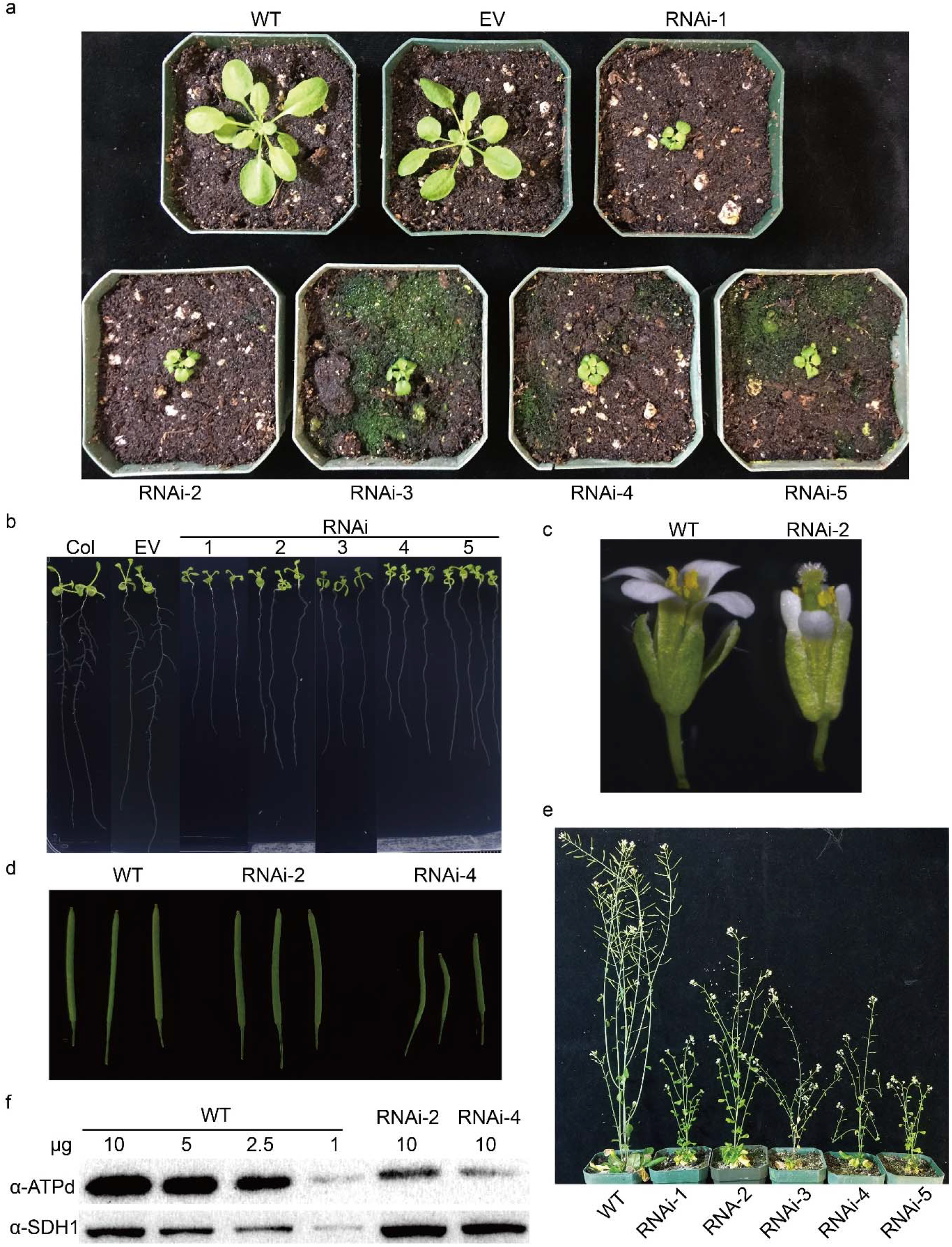
Knockdown of *ATPd* expression by RNAi affects plant growth, flower development and fertility. (a) Wild type Col-0, empty vector control (EV) and 5 *ATPd* RNAi lines grown for 4 weeks on soil in long day (16h light / 8h dark) growth conditions. (b) Phenotype of 12-day old seedlings grown on agar media. Knockdown plants showed shorter root length compared to the wild type and EV. (c) Effects of *ATPd* knockdown on flower morphology. A flower from RNAi line 2 showed underdeveloped petals and stamens. (d) Effects of *ATPd* knockdown on siliques. RNAi lines show shorter, deformed siliques. A few siliques of RNAi-2 could develop as fully as the wild type, but none of the siliques from the other RNAi lines could. (e) Wild type and RNAi plants grown for 74 days and 86 days, respectively, on soil in a long day (16h light / 8h dark) growth condition. (f) An immunoblot with anti-ATPd antibody showing the effectiveness of RNAi knockdown in two independent lines. Protein extracts from purified mitochondria of wild type and two *ATPd* RNAi plants were resolved by SDS–PAGE and probed with antibodies against ATPd and SDH1 (succinate dehydrogenase 1, loading control). The experiments were repeated three times with similar results.

We next performed immunoblot and qRT-PCR analysis to check the effectiveness of RNAi knockdown at both the protein and RNA level. As expected, all RNAi lines showed reduced levels of ATPd accumulation at both RNA and protein levels (Figure S2c and S2d). Immunoblot of ATPd with total seedling protein extracts showed a weak signal and a non-specific band right above subunit d preventing accurate quantification. To increase sensitivity of immunoblot signals, we repeated the experiment with purified mitochondria from two representative RNAi lines, lines 2 and 4. Immunoblots with mitochondrial proteins resulted in a clear signal without the non-specific band. The two RNAi lines accumulate subunit d at 10-25% of the wild-type level confirming the effectiveness of the RNAi knock-down (Figure 3f). *RNAi-2* plants showed slightly higher expression of ATPd than *RNAi-4* plants, which is consistent with an overall milder phenotype (Figure 3). The transcript levels of nuclear encoded ATP synthase subunit β and δ were not significantly different from the wild type (Figure S2d), indicating that knockdown of *ATPd* did not affect expression of other ATP synthase subunits at the transcriptional level.

### ATPd is essential for the assembly of holoenzyme, but does not affect other OXPHOS complexes

To determine the extent to which a decrease in ATP synthase subunit d affects the assembly of the ATP synthase complex or other OXPHOS complexes, we performed Blue Native-PAGE (BN-PAGE) analysis with purified mitochondrial proteins. The overall banding pattern of the Coomassie blue–stained gel looked similar for the RNAi lines and wild type with respect to the amount and sizes of super complexes I+III, complex I and complex III (Figure 4a). However, the band corresponding to the F_1_F_O_ complex was significantly decreased in RNAi lines. ATPase hydrolysis, as assessed by an in-gel activity assay, confirmed reduced activity of complex V (Figure 4b). These results indicate that subunit d is important for the stability and/or assembly of ATP synthase. Complex I and IV in-gel activity staining assays were also performed to determine if destabilization of the ATP synthase complex affects other OXPHOS complexes. We did not see differences between the wild type and RNAi lines for either complex I or complex IV activity (Figure 4c and d). In humans, subunit d is essential for the assembly of ATP synthase (Fujikawa *et al.*, 2015; He *et al.*, 2020). We conclude that ATP synthase subunit d is essential for the assembly of the F_1_F_O_ holoenzyme in *A. thaliana*, consistent with results from analysis of human mitochondria.

**Figure 4.**
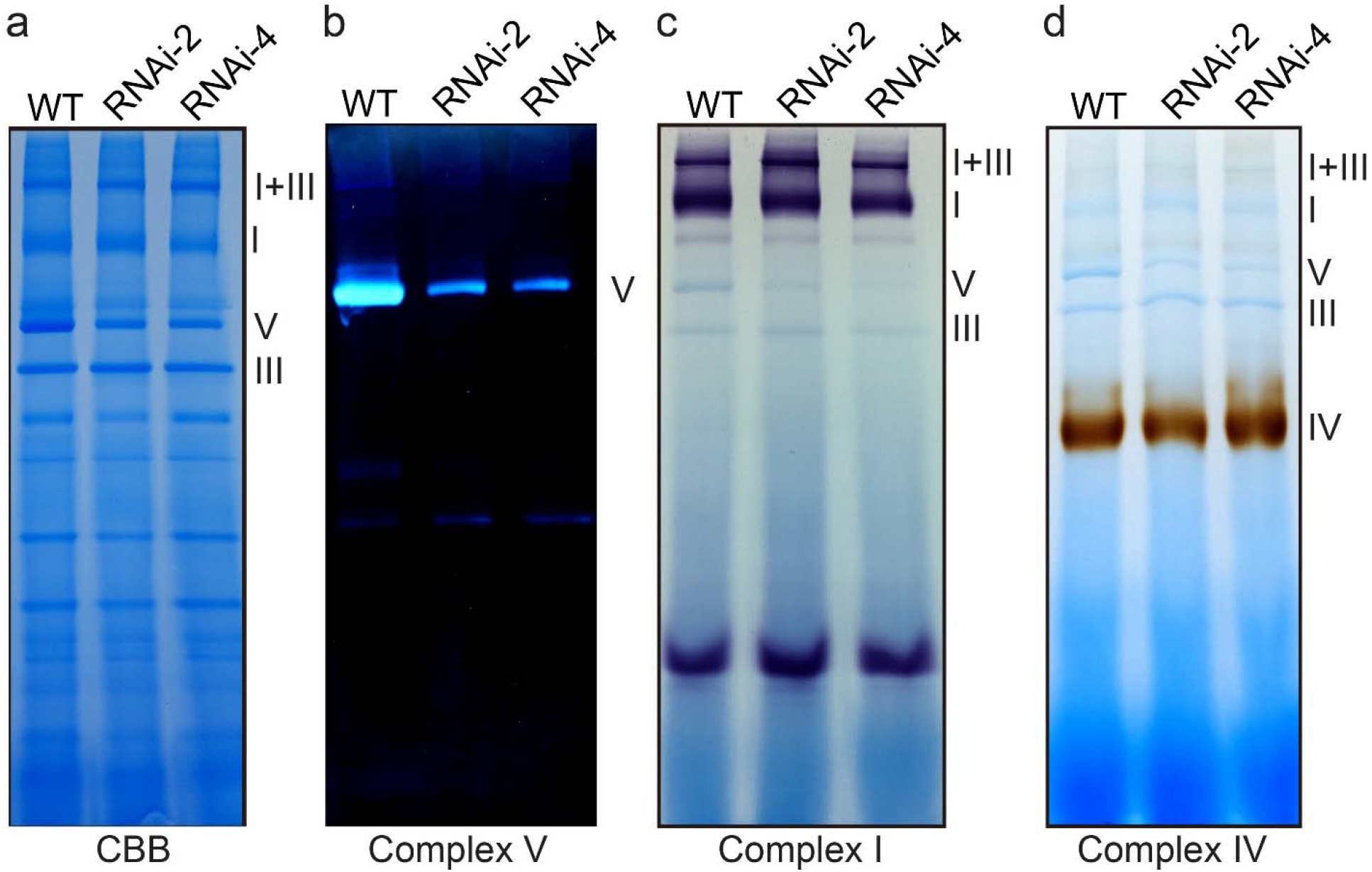
Knockdown of *ATPd* reduces the accumulation of complex V, but does not affect other OXPHOS complexes. BN-PAGE analysis of digitonin-solubilized mitochondria prepared from the wild type and two *ATPd* RNAi lines grown in half strength MS liquid media supplemented with 0.5 % sucrose. Gels were Coomassie-stained (a) or stained for ATPase hydrolysis (complex V, b) or NADH oxidase (complex I, c) or cytochrome c oxidase activity (complex IV, d) as indicated. The Coomassie staining of BN-PAGE gels was repeated four times with similar results. Activity staining for complex I, IV and V was repeated once, three times and twice, respectively. Similar results were obtained for complex IV and complex V activity staining.

To investigate in more detail how reduced ATP synthase subunit d could impact homeostasis of mitochondrial proteins, we examined the total proteome of mitochondria from *RNAi-2* and wild type using label-free quantitative proteomics. *RNAi-2* was chosen because it showed mildest phenotypes producing enough seeds for the experiment. A total of 1652 protein groups were detected from all samples (Figure 5a and Dataset S1). A majority of the ATP synthase subunits were significantly downregulated in *RNAi-2* except for subunits a, ε, e, g and IF_1_ (Figure 5a and b), indicating that most unassembled subunits are probably being degraded. There was no significant difference in subunits of any of the other OXPHOS complexes (Figure 5a). These data parallel the results obtained by BN-PAGE (Figure 4). Enrichment analysis of the proteomics data indicated that proteins involved in KEGG pathways like glutathione metabolism, sulfur metabolism, starch and sucrose metabolism, phagosome 2-oxocarboxylic acid metabolism, biosynthesis of amino acids, TCA cycle and glycolysis/gluconeogenesis are less abundant in *RNAi-2* (Figure S3 and Table S1). Overall downregulation of the proteins in these pathways reflect metabolic adjustment upon reduced complex V, which was also observed in a previous study (Geisler *et al.*, 2012).

**Figure 5.**
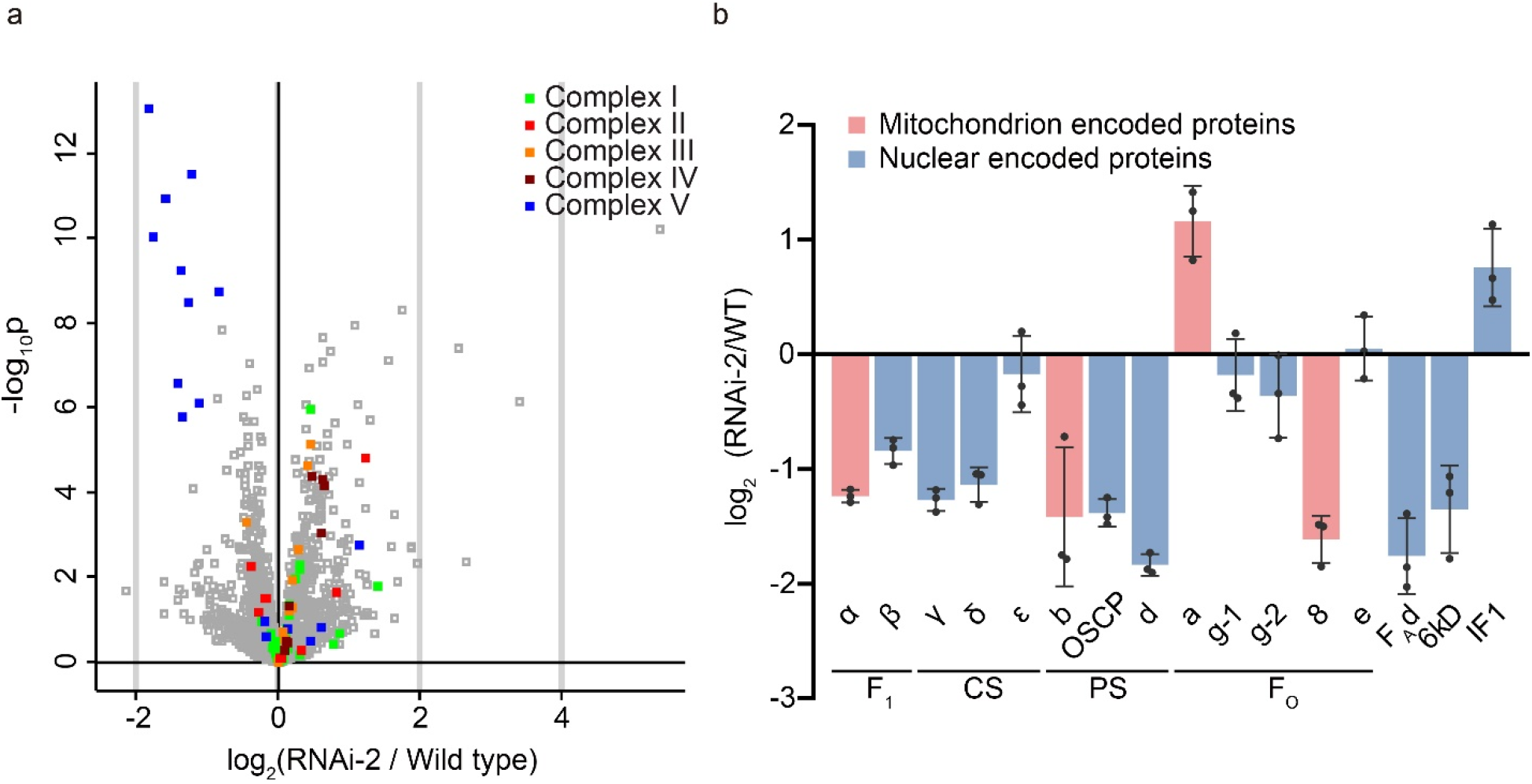
Most subunits of mitochondrial ATP synthase are downregulated in an *ATPd* RNAi knockdown line. (a) A volcano plot of the 1652 proteins identified in the mitochondrial proteomics experiment is shown. Subunits of each OXPHOS complex are colored differently. (b) Relative abundance of complex V subunits in RNAi-2 compared to wild type is shown. F_1_, CS, PS, F_O_ indicate F_1_ subcomplex, central stalk, peripheral stalk and F_O_ subcomplex of ATP synthase, respectively. Error bars represent SD of 3 data points (three biological replicates). The data for the identified proteins are given in Dataset S1.

### ATPd knockdown triggers mitochondrial retrograde signaling

MDS genes are known to be activated upon ROS accumulation caused by short-term inhibition of OXPHOS complexes by chemical inhibitors, as well as upon constitutive mitochondrial dysfunction resulting from genetic mutations (Clifton *et al.*, 2005; Schwarzländer *et al.*, 2012; De Clercq *et al.*, 2013; Van Aken *et al.*, 2016; Shapiguzov *et al.*, 2019). Therefore, we used qRT-PCR to investigate whether the constitutive inhibition of ATP synthase by knockdown of subunit d would induce MDS genes. Among about 30 MDS genes reported (De Clercq *et al.*, 2013), we selected genes whose protein products are targeted to mitochondria (*AOX1s*, *UPOX1*, *MGE1*, *NDB4*, *OM66*, *HSP23.5*), along with *ANAC013,* which encodes an ER membrane-localized transcription factor that can be transported to the nucleus upon sensing mitochondrial ROS, and *SOT12* which encodes sulfotransferase12 involved in cytosolic sulfate assimilation (Bohrer *et al.*, 2015). We additionally included *HSP23.6*, a close paralog of *HSP23.5*, because we found a mitochondrion dysfunction motif (MDM) (De Clercq *et al.*, 2013) in the promoter region (Figure S7). As expected, expression of most MDS genes was significantly increased in *RNAi-4* line (Figure 6), but more mildly in the *RNAi-2* line, which has a less severe phenotype and higher residual subunit d (Figure 3f) and complex V activity than *RNAi-4* (Figure 4b). Among six mitochondria targeted MDS proteins, most of them (AOX1a (2.91-fold), AOX1c (5.77-fold), MGE1 (1.64-fold), HSP23.5 (41.88-fold) and HSP23.6 (3.19-fold)) except OM66 (no change) were increased in our proteomics experiment using *RNAi-2* line (Dataset S1), showing high correlation with the qRT-PCR data. In conclusion, these results suggest that, similarly to acute inhibition of complex V with chemical inhibitors such as oligomycin, constitutive reduction of complex V activity by RNAi knockdown of ATPd can also trigger MRR.

**Figure 6.**
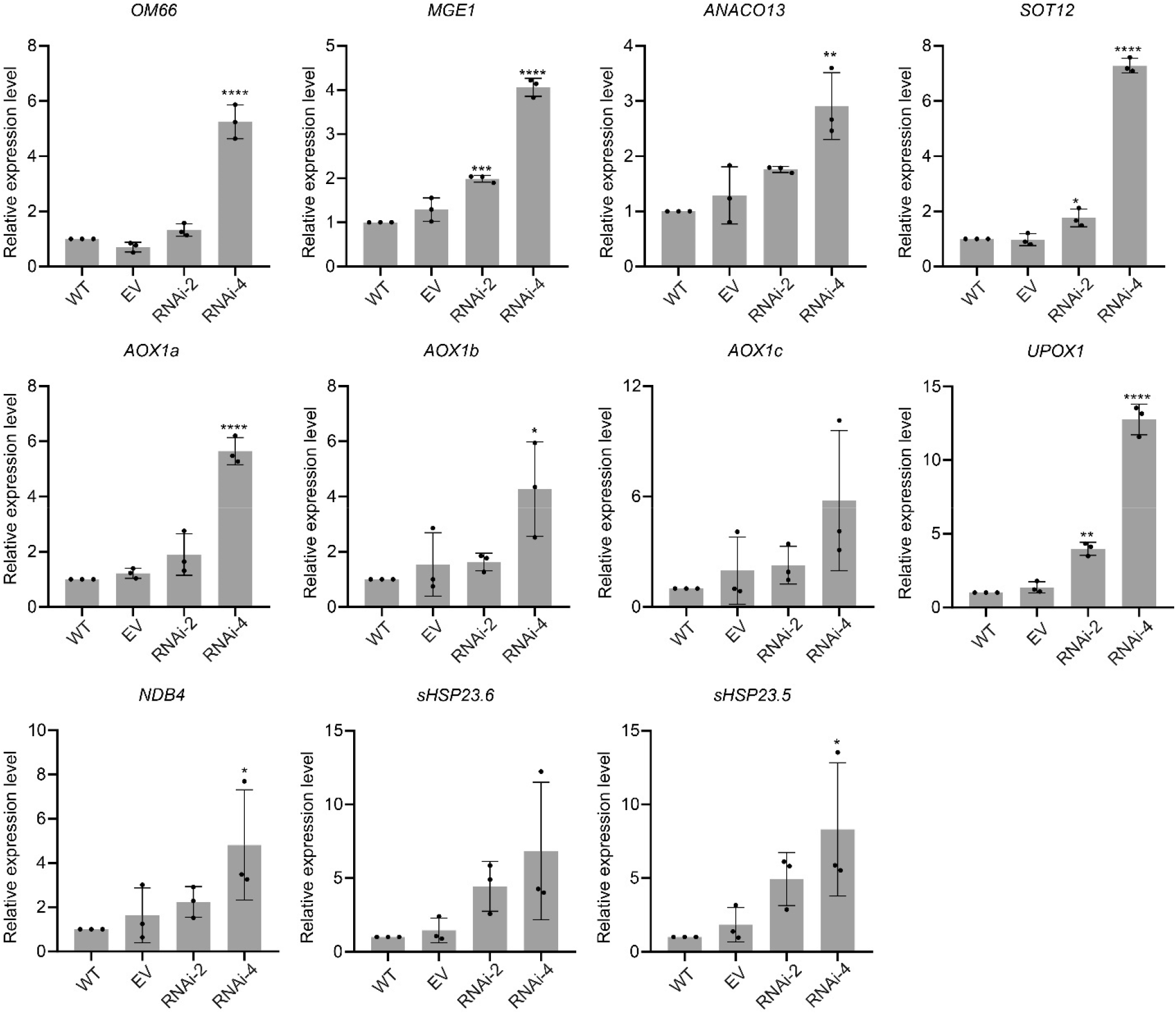
Transcript abundance of MDS genes from two RNAi lines were compared with the wild type (WT) and the empty vector control (EV) by qRT-PCR. Error bars are SD based on three biological replicates. *: p value<0.05, **: p value<0.01, ***: p value<0.001, ****: p value<0.0001 by one-way ANOVA.

### *ATPd* RNAi lines are more sensitive to heat stress

Mitochondrial mutants with defects in OXPHOS complexes are known to be affected in various abiotic stress responses. Notably, the *shot1-2* mutant, which is defective in mitochondria-targeted mTERF18 (mitochondria transcription termination factor 18), has reduced ATP synthase subunit d and is more tolerant to heat stress (Kim *et al.*, 2020). To investigate the effects of a defective ATP synthase complex on heat stress tolerance, we subjected the *ATPd* RNAi lines to heat stress treatment. 2.5-day old dark-grown seedlings were exposed to heat treatment at 45 °C for 3 hours after an acclimation treatment at 38 °C for 1.5 h followed by 22 °C for 2 h. Another set of seedlings was grown in parallel as a control for measuring growth without heat treatment. Hypocotyl elongation was measured 2.5 days after the treatment and compared with seedlings of the wild type, the empty vector (EV) control, and heat-sensitive *hot1-3* mutant, which is null for the cytosolic molecular chaperone HSP101 (Hong and Vierling, 2000). Without heat treatment, there were no significant growth delays observed with all five RNAi lines compared to wild type (Figure 7a), contrary to growth defects observed at a later developmental stage (Figure 3a). These data indicate that the function of the ATP synthase is not critical at this early stage of plant development when seedlings are grown on minimal nutrient medium. After the heat treatment, the growth of RNAi lines was more inhibited than the wild type and the EV control, although to a lesser degree than *hot1-3,* which shows essentially no growth after this treatment (Figure 7b). 10-day old light-grown seedlings grown on plates were also exposed to heat treatment. At this stage without heat treatment, the RNAi lines were smaller than wild type, but otherwise appear vigorous (Figure S3). After heat treatment, the growth of RNAi plants was also more inhibited than the wild type, with severely reduced growth and more dead seedlings (Figure 7c). Taken together, the heat sensitivity of the RNAi plants suggests that the function of ATP synthase is necessary for full recovery of plants after heat stress.

**Figure 7.**
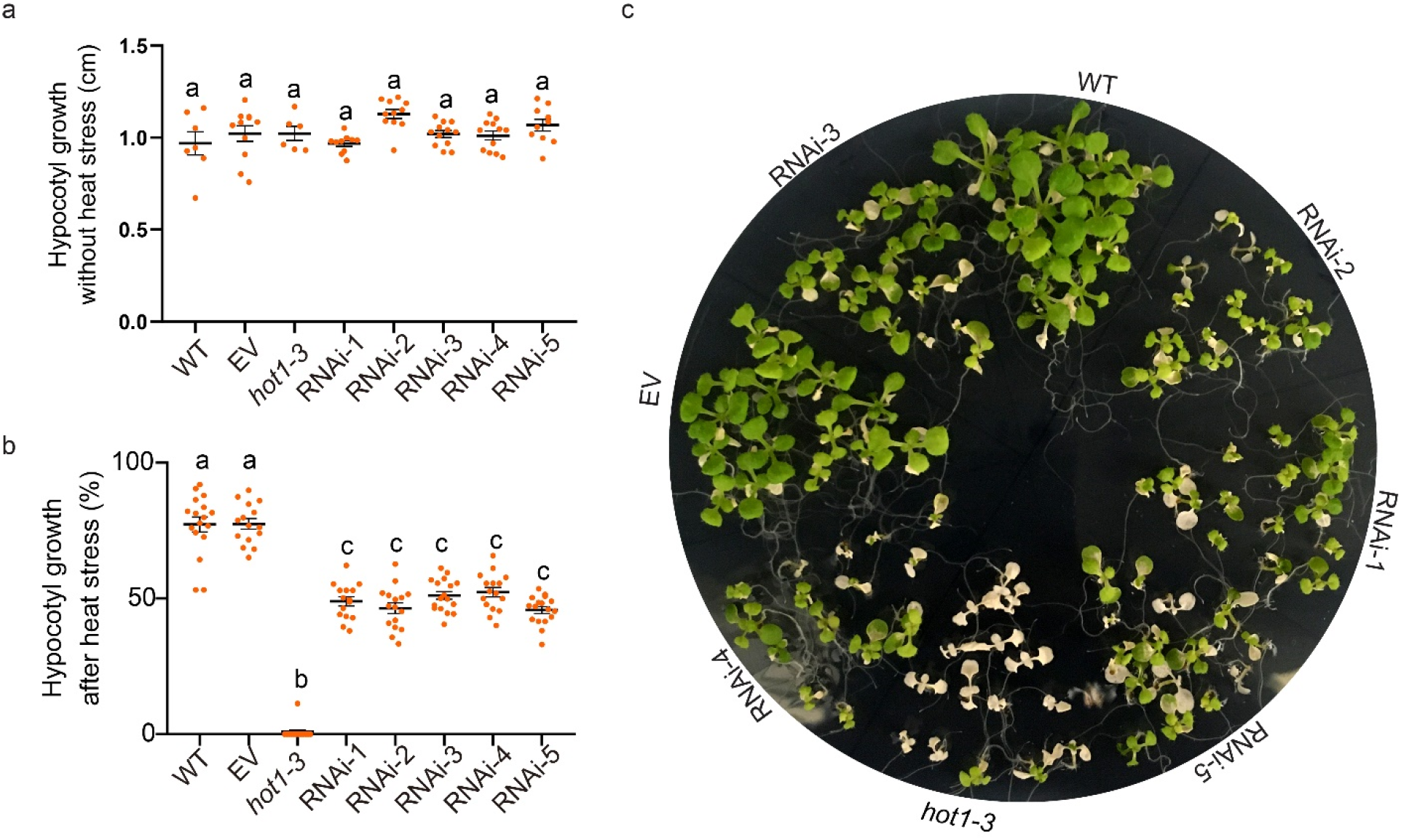
*ATPd* RNAi lines are more sensitive to heat stress. (a) Hypocotyl elongation of dark-grown seedlings grown on half strength MS agar media from 2.5 days to 5 days without heat stress. Error bars indicate SD; n ≥ 7. (b) Hypocotyl elongation assay shows more sensitivity of *ATPd* RNAi lines to heat stress treatment. Seedlings were grown on plates in the dark for 2.5 d and heat acclimated at 38°C for 90 min followed by 2 h at 22°C, then heat-shocked at 45°C for 3 h. Hypocotyl elongation 2.5 d after heat treatment was measured and expressed as a percentage of growth during the same time period for seedlings that had not been heat treated. Error bars indicate SD; n ≥ 16. Different letters on top of each column represent the significance at the 0.0001 level by one-way ANOVA. (c) Ten-day-old seedlings were heat-treated at 45°C for 3h after acclimation. Photograph was taken 7 days after heat treatment. The heat stress assay was repeated 6 time with similar results.

Heat shock proteins (HSPs) play a major role in heat stress tolerance. To investigate if decreased thermotolerance of the RNAi lines is due to altered HSP expression levels, we performed immunoblot analysis using antibodies against HSP101, cytosolic HSP70, sHSP class I, sHSP class II and mitochondrial HSP26.5 on total proteins from RNAi lines 2 and 4 either before or after heat stress. However, no significant difference between wild type and RNAi lines was observed in the protein levels of any of these HSPs (Figure S5).

### *ATPd* RNAi lines accumulate more hydrogen peroxide

As mitochondrial electron transport is a major source of ROS and altered ROS accumulation influences tolerance to abiotic stresses (Møller, 2001; Dutilleul *et al.*, 2003; Noctor *et al.*, 2007; Van Aken *et al.*, 2009; Rasmusson and Møller, 2011; Jacoby *et al.*, 2012; Waszczak *et al.*, 2018), we examined accumulation of H_2_O_2_, a major component of ROS, by DAB (3,3‘-diaminobenzidine) staining. 14-day old seedlings of wild type, *shot1-2* (as a control for reduced ROS (Kim *et al.*, 2012)) and two RNAi lines were stained with DAB after heat stress with or without acclimation. Wild-type plants after 30 minutes of heat stress at 45 °C accumulated more H_2_O_2_ than those kept at room temperature, while acclimation treatment reduced the ROS level. However, RNAi lines accumulated more H_2_O_2_ in all conditions, while *shot1-2* accumulated less ROS compared to wild type (Figure 8), indicating that decreased activity of ATP synthase negatively affects normal flow of electrons in the electron transport chain, constitutively producing more ROS in the RNAi lines. These results also suggest that the heat sensitivity of the RNAi plants could be attributed in part to increased ROS and likely oxidative damage associated with heat stress.

**Figure 8.**
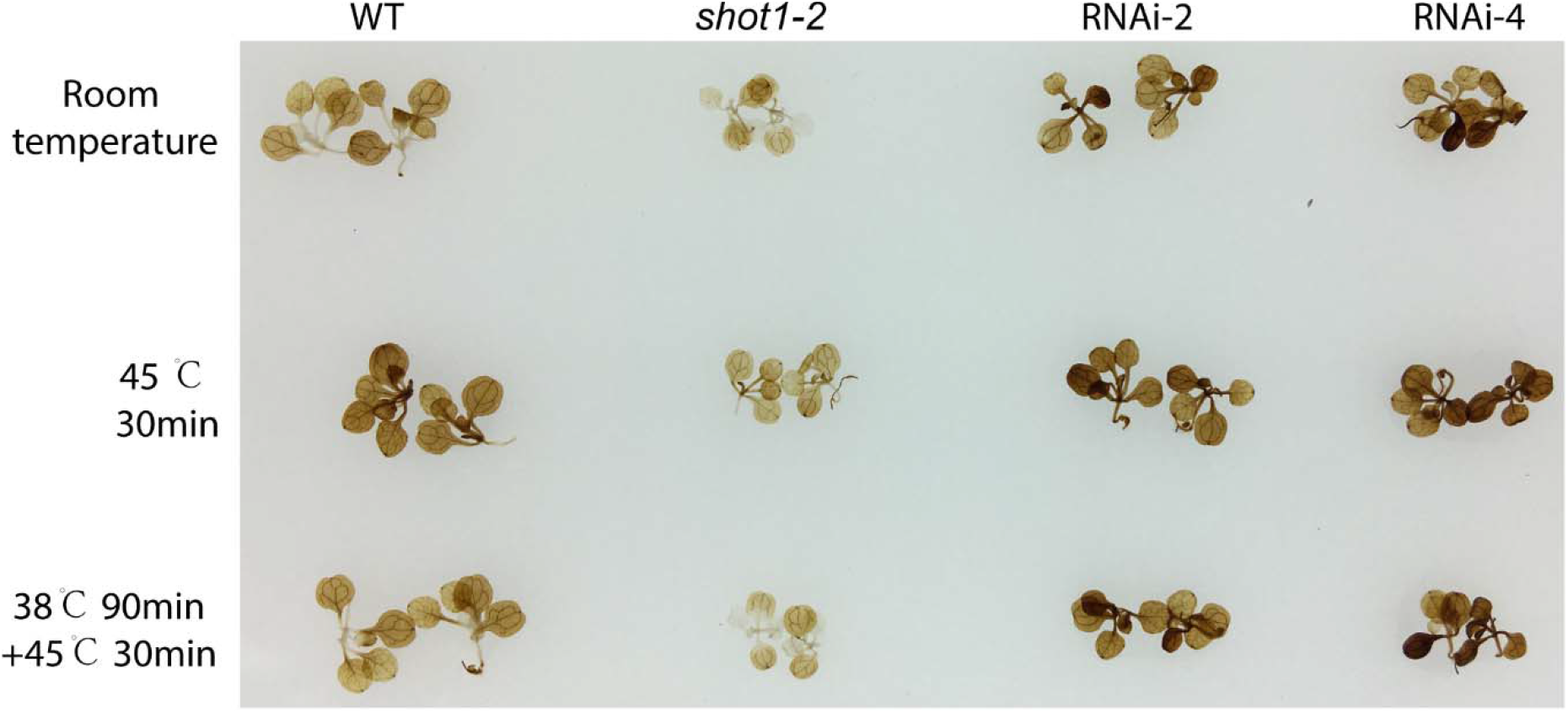
*ATPd* RNAi lines accumulated more hydrogen peroxide. 14-day old plants were stained with 3,3‘-diaminobenzidine (DAB). Heat treatments at 45°C for 30 min with or without acclimation (38 °C, 90 min followed by 2 h at room temperature) were performed before DAB staining. The experiments were repeated four times with similar results.

To investigate if knocking down ATPd affects tolerance to other stresses, sensitivity tests to oxidative, salt and drought stresses were carried out by assessing root growth on plates supplemented with paraquat (oxidizing agent), NaCl (salt stress) or PEG (polyethylene glycol, as a drought stress agent). However, no significant differences were found for all of these stresses between the RNAi lines and wild type (Figure S5), implying that low ATP synthase activity specifically affects thermotolerance under the conditions tested.

## DISCUSSION

We have demonstrated that subunit d belonging in the ATP synthase peripheral stalk is indispensable for ATP synthase assembly and function, and further confirmed the essential role of ATP synthase in pollen development and fertility. We isolated two different T-DNA insertion mutants in the *ATPd* gene encoding the subunit d of mitochondrial ATP synthase. While the *atpd-2* mutation with a T-DNA insertion in the last intron does not disrupt gene function, the *atpd-1* mutation with a T-DNA insertion in the second intron clearly disrupts the gene leading to lethality (Figure 1). The *atpd-1* mutation completely blocks pollen development at the tetrad stage, as well as severely reduces transmission efficiency (22 %) through the female gametophyte (Figure 2). It was previously shown that T-DNA knockout mutants of subunits δ and F_A_d also display male sterility and female defects analogous to *atpd-1* (Li *et al.*, 2010; Geisler *et al.*, 2012). Similar phenotypes are commonly observed in CMS mutants of other plant species with defects in ATP synthase subunits encoded in the mitochondrial genome (reviewed in Zancani *et al.*, 2020). A mitochondrial proteomics study in wheat revealed that decrease of ATP synthase subunits α, β and δ is related to CMS (Wang *et al.*, 2015). The *ATP6* gene (subunit a), one of the most rearranged genes in plant mitochondrial genomes, is associated with CMS in sunflower and pepper (Ji *et al.*, 2014, Makarenko *et al.*, 2019). The alteration of mRNA editing or gene rearrangements in *ATP9* (subunit c) lead to CMS in stem mustard, soybean, ramie, rice, tobacco and sunflower (Zancani *et al.*, 2020). In rice, the unedited *ORFB* (subunit 8, ATP8) transcript is responsible for male sterility (Das *et al.*, 2010). In agreement with CMS mutations in ATP synthase genes, our study strongly supports that, compared with other developmental processes, pollen development demands higher energy derived from mitochondrial OXPHOS, making it most vulnerable to mitochondrial dysfunction.

Since the *atpd-1* mutant was lethal, ATPd knockdown RNAi lines were generated to assess the effects of constitutively reduced levels of ATPd. Using stable knockdown lines allowed detailed phenotypic characterization of constitutively downregulated ATP synthase at different levels, which was difficult in previous studies using inducible knockdown of subunits OSCP, γ, and δ (Robison *et al.*, 2009; Geisler *et al.*, 2012). Subunit d protein levels of the five RNAi plants vary in a relatively small range between ~10 % and ~20 % of wild type (Figure 3f and S2c). Differences in the degree of growth retardation among the five RNAi lines were subtle, reflecting the small differences at the protein level. However, at the reproductive stage, the small differences at the protein level had significant impact on floral development leading to major differences in fertility. These results again highlight the critical role of complex V and mitochondrial function at the reproductive stage.

Dimers of mitochondrial ATP synthase oligomerize along the cristae edges playing critical roles in folding the mitochondrial inner membrane (Paumard *et al.*, 2002). Consistently, reduced cristae formation was observed in electron microscopy images of knockouts of subunits F_A_d and δ in *A. thaliana* (Li *et al.*, 2010; Geisler *et al.*, 2012). Therefore, it is expected that cristae formation is negatively affected in the ATPd RNAi lines. Cristae membranes are also the location of the other OXPHOS complexes, assembly of which could be dependent on proper cristae morphology. However, knockdown of the ATP synthase subunits only affected complex V biogenesis leaving the other OXPHOS complexes intact (Figure 4 and 5) suggesting that the biogenesis of ETC components is independent of complex V and its potentially negative effects on cristae membrane formation. Electron microscope images of mitochondria from the RNAi lines would be needed to fully support this conclusion.

The assembly of complex V in human mitochondria has been investigated recently using a CRISPR knockout of subunit d and F_6_ in HAP1 cell lines (He *et al.*, 2020). The PS (subunits b, d, F_6_ and OSCP in humans) is built up starting with subunit b by addition of subunits e and g together, followed by subunit f. This b-e-g-f complex is then assembled with a preformed F_1_-C_8_ module (α_3_, β_3_, γ, δ, ε, c_8_) by subunits OSCP and F_6_. Subunit d is the final component added to this intermediate complex before accepting the two subunits encoded in human mitochondrial genome, a and 8. This assembly pathway is supported by near normal accumulation of all subunits except subunits a and 8 in the subunit d CRISPR knockout (He *et al.*, 2020). Overall, the complex V assembly pathway is well conserved in yeast where defects in the assembly pathway also lead to a significant accumulation of assembly intermediates (Naumenko *et al.*, 2017). In contrast, our proteomics data show that all subunits except a, ε, e and g are downregulated in ATPd knockdown plants (Figure 5), indicating either a different assembly pathway for complex V in plants or extreme instability of assembly intermediates. In addition to subunits a and 8, *A. thaliana* has three more mitochondria-encoded subunits of complex V (α, b, and c). Plants also lack the F_6_ subunit of the human PS, but contain plant specific subunits F_A_d and 6 kDa whose function is not exactly defined (Zancani *et al.*, 2020). Even with these differences, it has been suggested that the assembly of plant complex V is similar to that in animals in that the matrix F_1_ intermediate is independently assembled before joining with the membrane-bound F_O_ domain (Li *et al.*, 2012; Meyer *et al.*, 2019). Our data, however, show no indication of significant accumulation of assembly intermediates, suggesting most likely that the assembly intermediates in plants are not as stable as those in yeast or human cell lines. Further research is required to elucidate the complex V assembly pathway in plants.

Acute perturbations of OXPHOS with chemical inhibitors such as antimycin A and oligomycin activate MRR, triggering induction of MDS genes in the nucleus (Schwarzländer *et al.*, 2012; De Clercq *et al.*, 2013). We demonstrated that reduction of ATPd by RNAi could also induce specific MDS genes (Figure 6). Expectedly, the strength of MRR signaling appears to be correlated with the degree of knockdown efficiency in RNAi plants. In *A. thaliana*, there are three mitochondria localized sHSPs, namely HSP23.5, HSP23.6 and HSP26.5, all of which are induced by heat stress (Waters and Vierling, 2020). We found that only HSP23.5 and HSP23.6 are induced by ATPd knockdown at both transcript and protein levels (Figure 6 and Dataset S1) while HSP26.5 is not detected (Dataset S1). The lack of MDM in the promoter sequence of HSP26.5 could explain the differential expression pattern of these sHSP genes. It suggests that the mitochondrial sHSPs may have diverged to perform different roles in responding to variable environmental stresses in higher plants.

ATP is essential for driving many biochemical reactions required not only for growth and development, but also for stress tolerance. Heat stress causes denaturation and aggregation of proteins, which need to be rescued by molecular chaperones such as HSP101 for recovery from heat damage in plants (Hong *et al.*, 2003). The majority of molecular chaperones essential for heat tolerance are dependent on ATP for their activity. Therefore, the heat sensitivity of ATPd knockdown plants could be explained by a reduced supply of ATP resulting from decreased complex V activity. Higher ROS in the ATPd knockdown plants likely also contributes to their heat sensitivity (Figure 7). Elevated ROS damages lipids, proteins, and nucleic acids. In ATPd knockdown plants, reduced activity of complex V could hyperpolarize the mitochondrial inner membrane and over-reduce ETC leading to increased leakage of electrons to O_2_ forming superoxide and other ROS. Increased activity of AOXs and NDBs in the ATPd knockdown plants could alleviate ROS formation, however, it does not appear to be sufficient for protection from stress. Enhanced heat sensitivity associated with increased ROS was also observed in other mitochondrial mutants such as the complex I-deficient *ndufs4* mutant (Kim *et al.*, 2012). In contrast, the *shot1-2* mutant with reduced accumulation of OXPHOS and ROS are more heat stress tolerant (Kim *et al.*, 2012; Kim *et al.*, 2020). Unexpectedly, the ATPd knockdown lines did not exhibit any significant sensitivity to other abiotic stress treatments (Figure S6), which increase ROS. Further studies of relationships between mitochondrial function, ROS and abiotic stress tolerance are required to understand the molecular mechanisms of abiotic stress tolerance involving mitochondria.

## EXPERIMENTAL PROCEDURES

### Plant materials and growth conditions

All *Arabidopsis thaliana* mutants are in the Col-0 background: *hot1-3* (Hong and Vierling, 2000), *qrt2* (Ogawa *et al.*, 2009), *shot1-2* (Kim *et al.*, 2012), *atpd-1* (SAIL_97_F05), and *atpd-2* (SAIL_515_E09). The T-DNA insertion mutants were provided by the Arabidopsis Biological Resource Center (ABRC). Seeds were surface sterilized and plated on half strength Murashige and Skoog (MS) medium (Sigma) containing 0.5% sucrose and 0.8% phytoagar or planted directly on soil. Plants were grown in growth chambers at 22°C/18°C, 16-h-day/8-h-night cycle or 12-h-day/night cycle with fluorescent lamps (50-80 μmol m^−2^ s^−1^). Seedlings for mitochondrial isolation were grown in 0.5x MS liquid media containing 0.5% sucrose with shaking in the same growth conditions. For thermotolerance assays, 2.5-d-old dark-grown and 10- to 12-d-old light-grown seedlings were treated as described previously (Hong and Vierling, 2000; Kim *et al.*, 2017).

### Generation of constructs and transgenic lines

The genomic *ATPd* DNA construct for complementation of the *atpd-1* mutation was generated by amplifying the appropriate DNA fragment (2438 bp) with the primers listed in Table S2 and cloning the fragment into the entry vector pCR8/GW/TOPO (Invitrogen). The fragment was inserted into the pMDC32 binary vector (Curtis and Grossniklaus, 2003) under control of the 35S promoter by Gateway LR cloning (Invitrogen). For generation of the RNAi knockdown plants, a 300 bp *ATPd* cDNA fragment near the 3’-end was amplified (see Table S2 for primer sequences) and cloned into the pENTR/D-TOPO vector (Invitrogen), resulting in pMK118. Using pMK118 as a template, the *ATPd* fragment was amplified with primers containing recognition sequences for restriction enzymes. The PCR fragment and vector pFGC5941 (ABRC stock # CD3-447) were digested with AscI and SwaI and ligated together to make pMK124. The same PCR fragment and pMK124 were digested with BamHI and XbaI and ligated together to make pMK127 (*ATPd* RNAi construct). The plasmid constructs were transformed into *Agrobacterium tumefaciens* GV3101. Transformation of *A. thaliana* was done by floral dipping (Clough and Bent, 1998). Transgenic plants were selected on media containing Hygromycin B (for the complementation construct) or Basta (for RNAi construct). Single insertion lines were selected by 3:1 segregation on selection media. For RNAi plants, the experiments were carried out with homozygous T3-T5 seeds.

### RT-PCR analysis

Total RNA for RT-PCR analysis was extracted from 10-12 day-old, plate-grown seedlings using TRIzol reagent (Invitrogen) or Plant RNA Isolation Kit (Beibei biotechnology) following the manufacturer’s instructions. 1-2μg of total RNA was reverse transcribed using the iScript gDNA Clear cDNA Synthesis kit (BioRad) or StarScript II First-strand cDNA Synthesis Kit (GeneStar). RT-PCR analysis was performed with gene-specific primers (Table S2). Primers of MGE1, ATPd, ATPδ, ATPβ and sHSP23.6 were designed with Primer Premier 5 program. Primers of OM66, ANAC013, SOT12, AOX1a, AOX1b, AOX1c, UPOX1, NBD4, sHSP23.5 and ACT were previously described (Clifton *et al.*, 2005; Geisler *et al.*, 2012; Kerchev *et al.*, 2014; Zhang *et al.*, 2014; Wu *et al.*, 2019; Adamowicz-Skrzypkowska *et al.*, 2020). qRT-PCR was performed with 2 × RealStar Green Power Mixture (gene star) in a QuantStudio 3 Real-Time PCR System (Applied Biosystems). Three biological replicates with three technical replicates were run for all genes. Data were analyzed with Microsoft Excel and GraphPad Prism 8.

### Isolation of mitochondria

Surface sterilized seeds (~20 mg) were grown in 50 ml of 0.5x MS liquid media containing 0.5% sucrose until stages 1.02-1.04 (Boyes *et al.*, 2001) in a growth chamber at 22°C/18°C, 16-h-day/8-h-night cycle with fluorescent lamps (60-80 μmol m^−2^ s^−1^). 15-30 grams of whole seedlings were harvested and ~40 ml extraction buffer (0.3 M sucrose, 25 mM K_4_P_2_O_7_, 10 mM KH_2_PO_4_, 2 mM EDTA, 1% [w/v] PVP-40, 1% [w/v] BSA, 5 mM cysteine, and 25 mM sodium ascorbate, pH7.5) per 10 grams of plant material was used to grind samples at 4°C using a mortar and pestle. The homogenate was spun at 1700g for 5 min and the resulting supernatant was then spun at 20000 x *g* for 10 min. The pellet was resuspended in wash buffer (0.3 M sucrose, 1 mM EGTA, and 10 mM MOPS/KOH, pH 7.2) and subjected to additional low (2500 x *g*)- and high (20000 x *g*)-speed centrifugations. The pellet was resuspended in a small volume (around 1 ml) of wash buffer and loaded onto a 0-4.4% PVP-40 gradient in 28% Percoll, 0.3 M sucrose, 10 mM MOPS/KOH, pH7.2. The gradient was centrifuged at 40000 x *g* for 45 min with SS-34 rotor in Sorvall RC 6 plus centrifuge (ThermoScientific). The mitochondrial fraction toward the bottom of the gradient was collected and washed three times by centrifugation at 30000 x *g* for 10 min. The final mitochondrial pellet was resuspended in ~1 ml wash buffer. The total protein concentration in the purified mitochondrial fraction was quantified by the Bradford assay and saved as a mitochondrial suspension at −80°C. Usually about 1 mg mitochondrial protein was obtained from about 30 g of fresh seedlings.

### SDS-PAGE and immunoblot analysis

SDS-PAGE and immunoblot analysis were performed as described previously (Kim *et al.*, 2012). Antibody dilutions were 1:1000 for antibodies against ATP synthase subunit d (PhytoAB 0593), HSP26.5 (Cocalico Biologicals) and cytosolic small HSPs (HSP17.6C-I and II) (Wehmeyer *et al.*, 1996); 1:2000 for anti-SDH (PhytoAB 0558); 1:5000 for antibodies against HSP101 (Hong and Vierling, 2000) and cytosolic HSP70 (Agrisera AS08 371); 1:5000 for donkey anti-rabbit IgG horseradish peroxidase conjugated (Amersham NA934-1ML). Pierce ECL Western Blotting Substrate (Thermo Scientific 32109) was used to visualize the signals with a G:Box iChemi XT(Syngene). Immunoblot experiments were repeated at least three times with similar results.

### Blue Native PAGE (BN-PAGE) and in-gel activity staining

Mitochondrial proteins (50 μg for Coomassie blue staining and 30μg for in-gel staining of complex activity) were solubilized (10 μg digitonin per μg of protein) in digitonin solution (5% digitonin, 30 mM HEPES, 150 mM potassium acetate, 10% glycerol) and prepared for electrophoresis as described (Eubel *et al.*, 2004). Protein complexes were separated in a 4.5 to 16% gradient gel. Following electrophoresis, gels were Coomassie-stained or subjected to ingel activity staining. In-gel activity staining for NADH oxidase (complex I) and ATP synthase (complex V) was carried out as described (Sabar *et al.*, 2005; Li *et al.*, 2019). Activity staining for complex I, IV and V was carried out twice, three times and twice, respectively.

### Detection of hydrogen peroxide by DAB staining

Surface sterilized seeds were grown on 0.5x MS agar media as described above for 2 weeks. Seedlings were either left untreated or heat stressed at 45 °C for 30 minutes with or without prior acclimation treatment (38 °C for 1.5h followed by 22 °C for 2h). DAB staining was carried out according to (Daudi and O’Brien, 2012). Three replicates were set up for each condition. Five seedlings per replicate per genotype were incubated with 2 ml DAB staining solution overnight in a 12-well plate in the dark after heat treatment. Pictures were taken on a light box after bleaching. The experiments were repeated four times with similar results.

### Stress treatment assays

Paraquat tolerance: Surface sterilized seeds were grown vertically on 0.5x MS agar media as described above for 4 days and then transferred to media containing 0,10, 50, 100, 500 mM of paraquat. Root growth was measured with Image J at 3 days after transfer and compared with growth on plates without paraquat as described previously (Kim *et al.*, 2012). At least 14 seedlings were tested for each genotype and each condition.

#### Salt stress

Surface sterilized seeds were grown vertically on 0.5x MS agar media with 50mM, 100mM and 150mM of NaCl for 6 days and then root growth was measured with Image J and compared with growth on plates without NaCl. At least 10 seedlings were measured for each genotype and each condition.

#### Drought stress

Surface sterilized seeds were grown vertically on PEG-infused 0.5x MS agar media plates. PEG-infused plates were prepared as described (Verslues *et al.*, 2006) with 25% and 30.5% of PEG8000. Root length was measured with image J after 6 days of growth and compared with growth on regular 0.5x MS media plates. At least 10 seedlings were measured for each genotype and each condition.

The paraquat, salt, and drought stress experiments were repeated three times, once and twice, respectively. Data were analyzed with Microsoft Excel and GraphPad Prism 8.

### Mitochondrial Proteomics

Wild-type Col-0 and *ATPd RNAi-2* were grown in 0.5x MS liquid media supplemented with 0.5% sucrose until stages 1.02-1.04 (Boyes *et al.*, 2001). 40 micrograms of mitochondrial proteins (three biological replicates) were separated by SDS-PAGE. The entire protein region of the gel was excised and subjected to in-gel trypsin (Promega V511A) digestion after reduction with DTT and alkylation with iodoacetamide. Peptides eluted from the gel were lyophilized and re-suspended in 25μL of 5% acetonitrile (0.1% (v/v) TFA). Downstream analyses were performed at the mass spectrometry facility at the University of Massachusetts Medical School. A 3μL injection (three technical replicates for each biological replicate) was separated on a Waters NanoAcquity UPLC in 5% acetonitrile (0.1% formic acid) at 4.0 μL/min for 4.0 min onto a 100 μm I.D. fused-silica pre-column packed with 2 cm of 5 μm (200Å) Magic C18AQ (Bruker-Michrom). Peptides were eluted at 300 nL/min from a 75 μm I.D. gravity-pulled analytical column packed with 25 cm of 3 μm (100Å) Magic C18AQ particles using a linear gradient from 5-35% of mobile phase B (acetonitrile + 0.1% formic acid) in mobile phase A (water + 0.1% formic acid). Ions were introduced by positive electrospray ionization via liquid junction at 1.5kV into a Thermo Scientific Q Exactive hybrid mass spectrometer. Mass spectra were acquired over m/z 300-1750 at 70,000 resolution (m/z 200) with an AGC (Automatic Gain Control) target of 1e6, and data-dependent acquisition selected the top 10 most abundant precursor ions for tandem mass spectrometry by HCD fragmentation using an isolation width of 1.6 Da. Peptides were fragmented by a normalized collisional energy of 27, and fragment spectra acquired at a resolution of 17,500 (m/z 200).

Raw data files were processed with MaxQuant (version 1.6.8.0) against the *A. thaliana* (Uniprot) FASTA file (downloaded 06/2019). Search parameters included Trypsin/P specificity, up to 2 missed cleavages, a fixed modification of carbamidomethyl cysteine, and variable modifications of oxidized methionine, and N-terminal acetylation. FDR was set to 1% at the protein and PSM level. Label free quantification was done using the MaxLFQ method (Cox *et al.*, 2014). The minimum number of unique peptides per protein group was 1. Unique and razor peptides were used for quantification. Downstream statistical analysis was performed using Perseus (version 1.6.12.0). Protein groups detected in at least five runs in either ATPq RNAi-2 or WT groups (9 runs in each group) were filtered resulting in 1652 total protein groups.

### DATA STATEMENT

The mass spectrometry proteomics data are openly accessible in MassIVE repository with the dataset identifier MSV000086794.

## ACCESSION NUMBERS

ATPα (ATMG01190), ATPβ (AT5G08670, AT5G08680, AT5G08690), ATPγ (AT2G33040), ATPδ (At5g47030), ATPε (AT1G51650), ATPa (ATMG00410), ATPb (ATMG00640), OSCP (AT5G13450), ATPd (AT3G52300), ATPe (AT5G15320), ATPg-1 (AT2G19680), ATPg-2 (AT4G29480), ATP8 (ATMG00480), F_A_d (AT2G21870), 6kDa (AT3G46430, AT5G59613), IF1 (AT2G27730), HSP23.5 (AT5G51440), HSP23.6 (AT4G25200), HSP26.5 (AT1G52560), OM66 (AT3G50930), ANAC013 (AT1G32870), SOT12 (AT2G03760), AOX1a (AT3G22370), AOX1b (AT3G22360), AOX1c (AT3G27620), UPOX1 (AT2G21640), NBD4 (AT2G20800), ACT (AT3G18780), HSP101 (AT1G74310), SHOT1 (AT3G60400), QRT2 (AT3G07970), MGE1 (AT5G55200)

## ACKNOWLEDGEMENTS

The authors acknowledge services of Light Microscopy Facility at the University of Massachusetts Amherst Institute of Applied Life Sciences Cores as well as Mass Spectrometry Facility at the University of Massachusetts Medical School for mitochondrial proteomics. Thanks for the support of China Scholarship Council to TX Liu. Work was supported by U.S. National Science Foundation grant IOS 1354960 to E.V.

## CONFLICT OF INTEREST

The authors declare no conflict of interest.

## SUPPORTING INFORMATION

**Figure S1**. Protein sequence alignment of ATP synthase subunit d homologs of *Arabidopsis thaliana* (Uniprot Q9FT52), *Oryza sativa* (Uniprot Q7XXS0), *Homo sapiens* (Uniprot O75947) and *Saccharomyces cerevisiae* (Uniprot P30902).

**Figure S2**. Phenotypes of *ATPd* RNAi knockdown plants.

**Figure S3**. Volcano plots of the 1652 proteins identified in the mitochondrial proteomics experiment are shown with colored squares representing proteins involved in specific categories of KEGG pathways.

**Figure S4**. Seedlings of wild type (WT), empty vector control (EV), *hot1-3* and five *ATPd* RNAi lines grown on half-strength MS agar media for 17 days without heat stress.

**Figure S5**. Protein levels of major cytosolic heat shock proteins (HSP) and a mitochondrial sHSP are not differentially regulated in *ATPd* RNAi lines.

**Figure S6**. No significant differences in paraquat, salt and drought stresses were observed between *ATPd* RNAi lines and wild type (WT).

**Figure S7**. Mitochondrial dysfunction motif (MDM) exists in the promoter region of sHSP23.5 and sHSP23.6 while not in sHSP26.5.

**Table S1**. Enrichment of different KEGG pathways from proteomics study comparing *ATPd* RNAi line with wild type.

**Table S2**. Primers used in this study.

**Dataset S1**. Proteomics dataset.

## Notes

### Competing Interest Statement

The authors have declared no competing interest.

## REFERENCES

Adamowicz-Skrzypkowska, A., Kwasniak-Owczarek, M., Van Aken, O., Kazmierczak, U. and Janska, H. (2020) Joint inhibition of mitochondrial complex IV and alternative oxidase by genetic or chemical means represses chloroplast transcription in Arabidopsis. Philos. Trans. R. Soc. B Biol. Sci., 375, 20190409.

Arnold, I., Pfeiffer, K., Neupert, W., Stuart, R.A. and Schagger, H. (1998) Yeast mitochondrial F1F0-ATP synthase exists as a dimer: identification of three dimer-specific subunits. Embo J., 17, 7170–7178.

Artika, I.M. (2019) Current understanding of structure, function and biogenesis of yeast mitochondrial ATP synthase. J. Bioenerg. Biomembr., 51, 315–328.

Blum, T.B., Hahn, A., Meier, T., Davies, K.M. and Kuhlbrandt, W. (2019) Dimers of mitochondrial ATP synthase induce membrane curvature and self-assemble into rows. Proc. Natl. Acad. Sci. U. S. A., 116, 4250–4255.

Bohrer, A.-S., Kopriva, S. and Takahashi, H. (2015) Plastid-cytosol partitioning and integration of metabolic pathways for APS/PAPS biosynthesis in Arabidopsis thaliana. Front. Plant Sci., 5. Available at: https://www.frontiersin.org/articles/10.3389/fpls.2014.00751/full [Accessed January 18, 2021].

Boyes, D.C., Zayed, A.M., Ascenzi, R., McCaskill, A.J., Hoffman, N.E., Davis, K.R. and Görlach, J. (2001) Growth Stage–Based Phenotypic Analysis of Arabidopsis: A Model for High Throughput Functional Genomics in Plants. Plant Cell, 13, 1499–1510.

Brugière, S., Kowalski, S., Ferro, M., et al. (2004) The hydrophobic proteome of mitochondrial membranes from Arabidopsis cell suspensions. Phytochemistry, 65, 1693–1707.

Chen, L. and Liu, Y.-G. (2014) Male Sterility and Fertility Restoration in Crops. Annu. Rev. Plant Biol., 65, 579–606.

Clifton, R., Lister, R., Parker, K.L., Sappl, P.G., Elhafez, D., Millar, A.H., Day, D.A. and Whelan, J. (2005) Stress-induced co-expression of alternative respiratory chain components in Arabidopsis thaliana. Plant Mol. Biol., 58, 193.

Clough, S.J. and Bent, A.F. (1998) Floral dip: a simplified method for Agrobacterium-mediated transformation of Arabidopsis thaliana. Plant J. Cell Mol. Biol., 16, 735–743.

Cox, J., Hein, M.Y., Luber, C.A., Paron, I., Nagaraj, N. and Mann, M. (2014) Accurate proteome-wide label-free quantification by delayed normalization and maximal peptide ratio extraction, termed MaxLFQ. Mol. Cell. Proteomics MCP, 13, 2513–2526.

Curtis, M.D. and Grossniklaus, U. (2003) A Gateway Cloning Vector Set for High-Throughput Functional Analysis of Genes in Planta. Plant Physiol., 133, 462–469.

Daudi, A. and O’Brien, J.A. (2012) Detection of Hydrogen Peroxide by DAB Staining in Arabidopsis Leaves. Bio-Protoc., 2, e263–e263.

De Clercq, I., Vermeirssen, V., Van Aken, O., et al. (2013) The Membrane-Bound NAC Transcription Factor ANAC013 Functions in Mitochondrial Retrograde Regulation of the Oxidative Stress Response in Arabidopsis. Plant Cell, 25, 3472–3490.

Dudkina, N.V., Oostergetel, G.T., Lewejohann, D., Braun, H.P. and Boekema, E.J. (2010) Row-like organization of ATP synthase in intact mitochondria determined by cryo-electron tomography. Biochim. Biophys. Acta-Bioenerg., 1797, 272–277.

Dutilleul, C., Garmier, M., Noctor, G., Mathieu, C., Chétrit, P., Foyer, C.H. and Paepe, R. de (2003) Leaf Mitochondria Modulate Whole Cell Redox Homeostasis, Set Antioxidant Capacity, and Determine Stress Resistance through Altered Signaling and Diurnal Regulation. Plant Cell, 15, 1212–1226.

Eubel, H., Heinemeyer, J. and Braun, H.-P. (2004) Identification and Characterization of Respirasomes in Potato Mitochondria. Plant Physiol., 134, 1450–1459.

Fujikawa, M., Sugawara, K., Tanabe, T. and Yoshida, M. (2015) Assembly of human mitochondrial ATP synthase through two separate intermediates, F _1_ - *c* -ring and *b* - *e* - *g* complex. FEBS Lett., 589, 2707–2712.

Geisler, D.A., Päpke, C., Obata, T., et al. (2012) Downregulation of the δ-Subunit Reduces Mitochondrial ATP Synthase Levels, Alters Respiration, and Restricts Growth and Gametophyte Development in Arabidopsis. Plant Cell, 24, 2792–2811.

He, J., Carroll, J., Ding, S., Fearnley, I.M., Montgomery, M.G. and Walker, J.E. (2020) Assembly of the peripheral stalk of ATP synthase in human mitochondria. Proc. Natl. Acad. Sci., 117, 29602–29608.

Hong, S.-W., Lee, U. and Vierling, E. (2003) Arabidopsis hot Mutants Define Multiple Functions Required for Acclimation to High Temperatures. Plant Physiol., 132, 757–767.

Hong, S.-W. and Vierling, E. (2000) Mutants of Arabidopsis thaliana defective in the acquisition of tolerance to high temperature stress. Proc. Natl. Acad. Sci. U. S. A., 97, 4392–4397.

Horn, R., Gupta, K.J. and Colombo, N. (2014) Mitochondrion role in molecular basis of cytoplasmic male sterility. Mitochondrion, 19, 198–205.

Jacoby, R.P., Li, L., Huang, S., Lee, C.P., Millar, A.H. and Taylor, N.L. (2012) Mitochondrial Composition, Function and Stress Response in Plants. J. Integr. Plant Biol., 54, 887–906.

Kerchev, P.I., De Clercq, I., Denecker, J., Mühlenbock, P., Kumpf, R., Nguyen, L., Audenaert, D., Dejonghe, W. and Van Breusegem, F. (2014) Mitochondrial Perturbation Negatively Affects Auxin Signaling. Mol. Plant, 7, 1138–1150.

Kim, M., Lee, U., Small, I., Francs-Small, C.C. des and Vierling, E. (2012) Mutations in an Arabidopsis Mitochondrial Transcription Termination Factor-Related Protein Enhance Thermotolerance in the Absence of the Major Molecular Chaperone HSP101. Plant Cell, 24, 3349–3365.

Kim, M., McLoughlin, F., Basha, E. and Vierling, E. (2017) Assessing Plant Tolerance to Acute Heat Stress. BIO-Protoc., 7. Available at: https://bio-protocol.org/e2405 [Accessed June 30, 2020].

Kim, M., Schulz, V., Brings, L., Schoeller, T., Kühn, K. and Vierling, E. (2020) Mitochondrial nucleoid organization and biogenesis of complex I require mTERF18/SHOT1 and ATAD3 in *Arabidopsis thaliana*. bioRxiv, 2020.05.11.088575.

Kühlbrandt, W. (2019) Structure and Mechanisms of F-Type ATP Synthases. Annu. Rev. Biochem., 88, 515–549.

Li, L., Carrie, C., Nelson, C., Whelan, J. and Millar, A.H. (2012) Accumulation of Newly Synthesized F1 in Vivo in Arabidopsis Mitochondria Provides Evidence for Modular Assembly of the Plant F1Fo ATP Synthase *. J. Biol. Chem., 287, 25749–25757.

Li, W.-Q., Zhang, X.-Q., Xia, C., Deng, Y. and Ye, D. (2010) MALE GAMETOPHYTE DEFECTIVE 1, Encoding the FAd Subunit of Mitochondrial F1F0-ATP Synthase, is Essential for Pollen Formation in Arabidopsis thaliana. Plant Cell Physiol., 51, 923–935.

Li, X.-L., Huang, W.-L., Yang, H.-H., Jiang, R.-C., Sun, F., Wang, H.-C., Zhao, J., Xu, C.-H. and Tan, B.-C. (2019) EMP18 functions in mitochondrial atp6 and cox2 transcript editing and is essential to seed development in maize. New Phytol., 221, 896–907.

Meyer, E.H., Welchen, E. and Carrie, C. (2019) Assembly of the Complexes of the Oxidative Phosphorylation System in Land Plant Mitochondria. Annu. Rev. Plant Biol., 70, 23–50.

Møller, I.M. (2001) PLANT MITOCHONDRIA AND OXIDATIVE STRESS: Electron Transport, NADPH Turnover, and Metabolism of Reactive Oxygen Species. Annu. Rev. Plant Physiol. Plant Mol. Biol., 52, 561–591.

Moore, R.C., Kozyreva, O., Lebel-Hardenack, S., Siroky, J., Hobza, R., Vyskot, B. and Grant, S.R. (2003) Genetic and Functional Analysis of DD44, a Sex-Linked Gene From the Dioecious Plant Silene latifolia, Provides Clues to Early Events in Sex Chromosome Evolution. Genetics, 163, 321–334.

Mühleip, A., McComas, S.E. and Amunts, A. (2019) Structure of a mitochondrial ATP synthase with bound native cardiolipin. Elife, 8, e51179.

Naumenko, N., Morgenstern, M., Rucktäschel, R., Warscheid, B. and Rehling, P. (2017) INA complex liaises the F 1 F o -ATP synthase membrane motor modules. Nat. Commun., 8, 1237.

Ng, S., Ivanova, A., Duncan, O., et al. (2013) A Membrane-Bound NAC Transcription Factor, ANAC017, Mediates Mitochondrial Retrograde Signaling in Arabidopsis. Plant Cell, 25, 3450–3471.

Noctor, G., De Paepe, R. and Foyer, C.H. (2007) Mitochondrial redox biology and homeostasis in plants. Trends Plant Sci., 12, 125–134.

Ogawa, M., Kay, P., Wilson, S. and Swain, S.M. (2009) ARABIDOPSIS DEHISCENCE ZONE POLYGALACTURONASE1 (ADPG1), ADPG2, and QUARTET2 Are Polygalacturonases Required for Cell Separation during Reproductive Development in Arabidopsis. Plant Cell, 21, 216–233.

Paumard, P., Vaillier, J., Coulary, B., Schaeffer, J., Soubannier, V., Mueller, D.M., Brèthes, D., Rago, J.-P. di and Velours, J. (2002) The ATP synthase is involved in generating mitochondrial cristae morphology. EMBO J., 21, 221–230.

Pinke, G., Zhou, L. and Sazanov, L.A. (2020) Cryo-EM structure of the entire mammalian F-type ATP synthase. Nat. Struct. Mol. Biol., 1–9.

Rasmusson, A.G. and Møller, I.M. (2011) Mitochondrial Electron Transport and Plant Stress. In F. Kempken, ed. Plant Mitochondria. Advances in Plant Biology. New York, NY: Springer, pp. 357–381. Available at: https://doi.org/10.1007/978-0-387-89781-3_14 [Accessed January 10, 2021].

Robison, M.M., Ling, X., Smid, M.P.L., Zarei, A. and Wolyn, D.J. (2009) Antisense Expression of Mitochondrial ATP Synthase Subunits OSCP (ATP5) and γ (ATP3) Alters Leaf Morphology, Metabolism and Gene Expression in Arabidopsis. Plant Cell Physiol., 50, 1840–1850.

Sabar, M., Balk, J. and Leaver, C.J. (2005) Histochemical staining and quantification of plant mitochondrial respiratory chain complexes using blue-native polyacrylamide gel electrophoresis. Plant J., 44, 893–901.

Schwarzländer, M., König, A.-C., Sweetlove, L.J. and Finkemeier, I. (2012) The impact of impaired mitochondrial function on retrograde signalling: a meta-analysis of transcriptomic responses. J. Exp. Bot., 63, 1735–1750.

Sessions, A., Burke, E., Presting, G., et al. (2002) A High-Throughput Arabidopsis Reverse Genetics System. Plant Cell, 14, 2985–2994.

Shapiguzov, A., Vainonen, J.P., Hunter, K., et al. (2019) Arabidopsis RCD1 coordinates chloroplast and mitochondrial functions through interaction with ANAC transcription factors. eLife, 8. Available at: https://elifesciences.org/articles/43284 [Accessed July 12, 2019].

Sun, X., Wheeler, C.T., Yolitz, J., Laslo, M., Alberico, T., Sun, Y., Song, Q. and Zou, S. (2014) A Mitochondrial ATP Synthase Subunit Interacts with TOR Signaling to Modulate Protein Homeostasis and Lifespan in Drosophila. Cell Rep., 8, 1781–1792.

Tan, Y.-F., Millar, A.H. and Taylor, N.L. (2012) Components of Mitochondrial Oxidative Phosphorylation Vary in Abundance Following Exposure to Cold and Chemical Stresses. J. Proteome Res., 11, 3860–3879.

Van Aken, O., Ford, E., Lister, R., Huang, S. and Millar, A.H. (2016) Retrograde signalling caused by heritable mitochondrial dysfunction is partially mediated by ANAC017 and improves plant performance. Plant J., 88, 542–558.

Van Aken, O., Zhang, B., Carrie, C., Uggalla, V., Paynter, E., Giraud, E. and Whelan, J. (2009) Defining the Mitochondrial Stress Response in Arabidopsis thaliana. Mol. Plant, 2, 1310–1324.

Verslues, P.E., Agarwal, M., Katiyar-Agarwal, S., Zhu, J. and Zhu, J.K. (2006) Methods and concepts in quantifying resistance to drought, salt and freezing, abiotic stresses that affect plant water status. (vol 45, pg 523, 2006). Plant J., 46, 1092–1092.

Wang, Y., Berkowitz, O., Selinski, J., Xu, Y., Hartmann, A. and Whelan, J. (2018) Stress responsive mitochondrial proteins in Arabidopsis thaliana. Free Radic. Biol. Med., 122, 28–39.

Waszczak, C., Carmody, M. and Kangasjärvi, J. (2018) Reactive Oxygen Species in Plant Signaling. Annu. Rev. Plant Biol., 69, 209–236.

Waters, E.R. and Vierling, E. (2020) Plant small heat shock proteins -evolutionary and functional diversity. New Phytol., 227, 24–37.

Welch, A.K., Bostwick, C.J. and Cain, B.D. (2011) Manipulations in the peripheral stalk of the Saccharomyces cerevisiae F1F0-ATP synthase. J Biol Chem, 286, 10155–62.

Wu, Z., Han, S., Zhou, H., Tuang, Z.K., Wang, Y., Jin, Y., Shi, H. and Yang, W. (2019) Cold stress activates disease resistance in Arabidopsis thaliana through a salicylic acid dependent pathway. Plant Cell Environ., 42, 2645–2663.

Zancani, M., Braidot, E., Filippi, A. and Lippe, G. (2020) Structural and functional properties of plant mitochondrial F-ATP synthase. Mitochondrion, 53, 178–193.

Zhang, B., Aken, O.V., Thatcher, L., et al. (2014) The mitochondrial outer membrane AAA ATPase AtOM66 affects cell death and pathogen resistance in Arabidopsis thaliana. Plant J., 80, 709–727.

