## Supplemental Figures and Tables for "Mitochondrial ATP Synthase Subunit d, a Component of the Peripheral Stalk, is Essential for Growth and Heat Stress Tolerance in *Arabidopsis thaliana*"

### Supporting figures

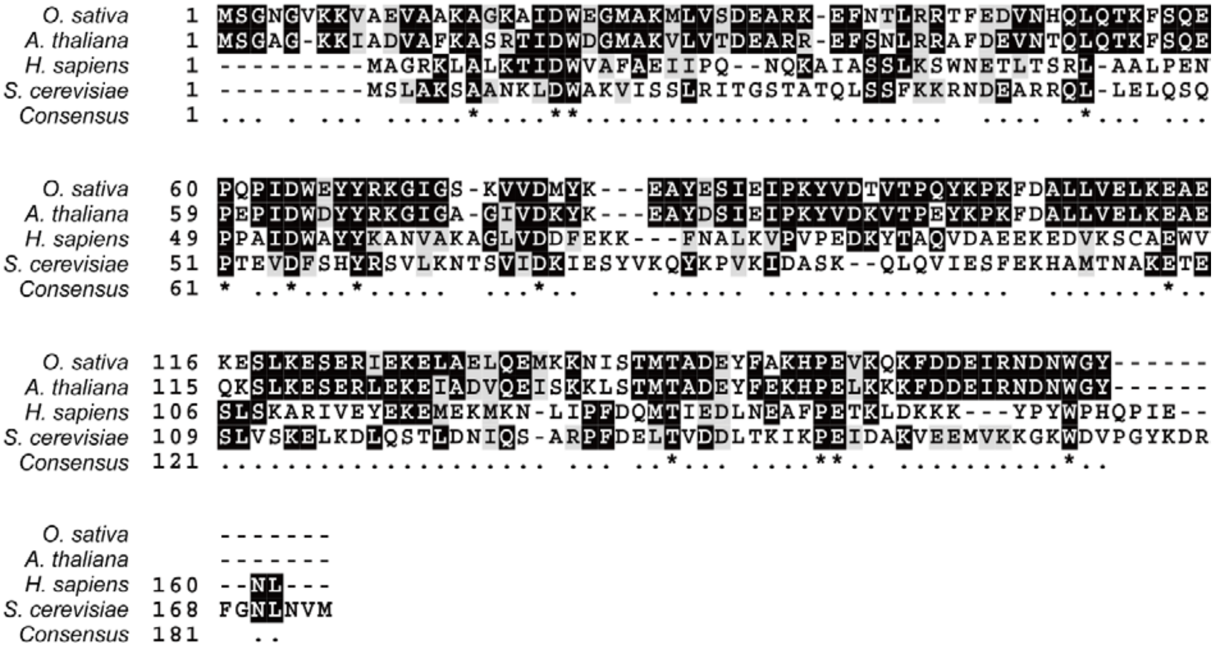


Figure S1. Protein sequence alignment of ATP synthase subunit d homologs of *Arabidopsis thaliana* (Uniprot Q9FT52), *Oryza sativa* (Uniprot Q7XXS0), *Homo sapiens* (Uniprot O75947) and *Saccharomyces cerevisiae* (Uniprot P30902). Identical residues are highlighted in black and similar residues in gray.

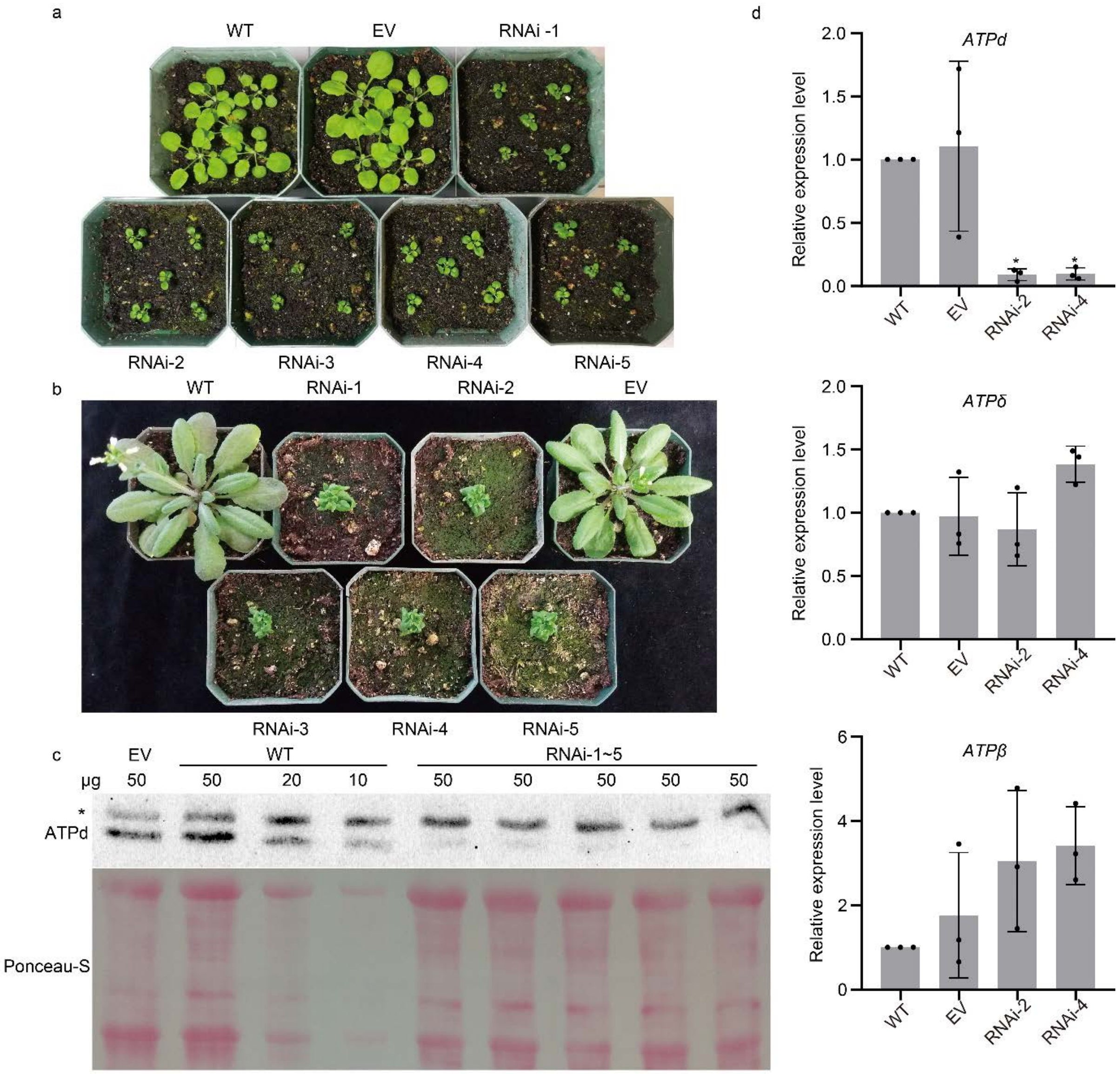


Figure S2. Phenotypes of *ATPd* RNAi knockdown plants. (a) Wild type (WT), Empty vector control (EV), and five *ATPd* RNAi lines were grown for 27 days on soil in long day (16h light / 8h dark) growth conditions. (b) Plants grown for 49 days as in (a). (c) ATPd protein levels in RNAi lines. Total proteins extracted from 12 d old seedlings of WT and five *ATPd* RNAi lines were resolved by SDS–PAGE and probed with antibody against ATPd. An asterisk indicates non-specific bands. The experiments were repeated more than three times with similar results. (d) Transcript abundance of subunits d (*ATPd*), δ (*ATPδ*) and β (*ATPβ*) of mitochondrial ATP synthase from two RNAi lines were compared to WT and EV by qRT-PCR. Error bars represent SD from three biological replicates. *: $P<0.05$ by one-way ANOVA.

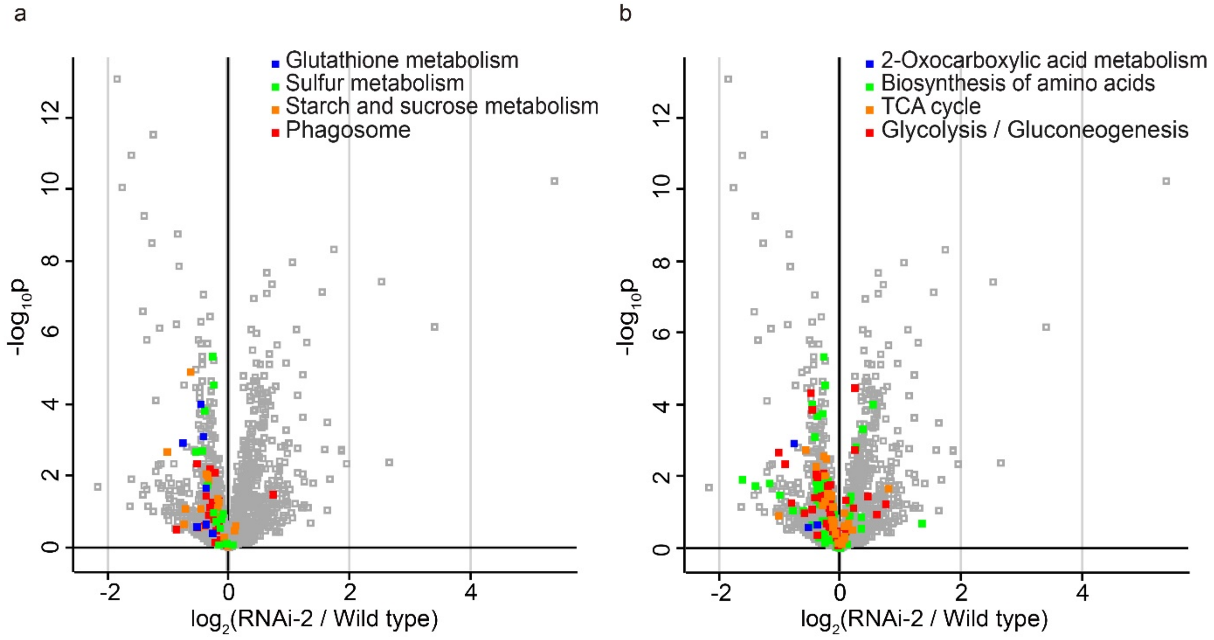


Figure S3. Volcano plots of the 1652 proteins identified in the mitochondrial proteomics experiment are shown with colored squares representing proteins involved in specific categories of KEGG pathways. (a) Proteins involved in glutathione metabolism, sulfur metabolism, starch and sucrose metabolism and phagosome are less abundant in *ATPd* RNAi line. (b) Proteins involved in 2-oxocarboxylic acid metabolism, biosynthesis of amino acids, TCA cycle and glycolysis/gluconeogenesis are less abundant in *ATPd* RNAi line.

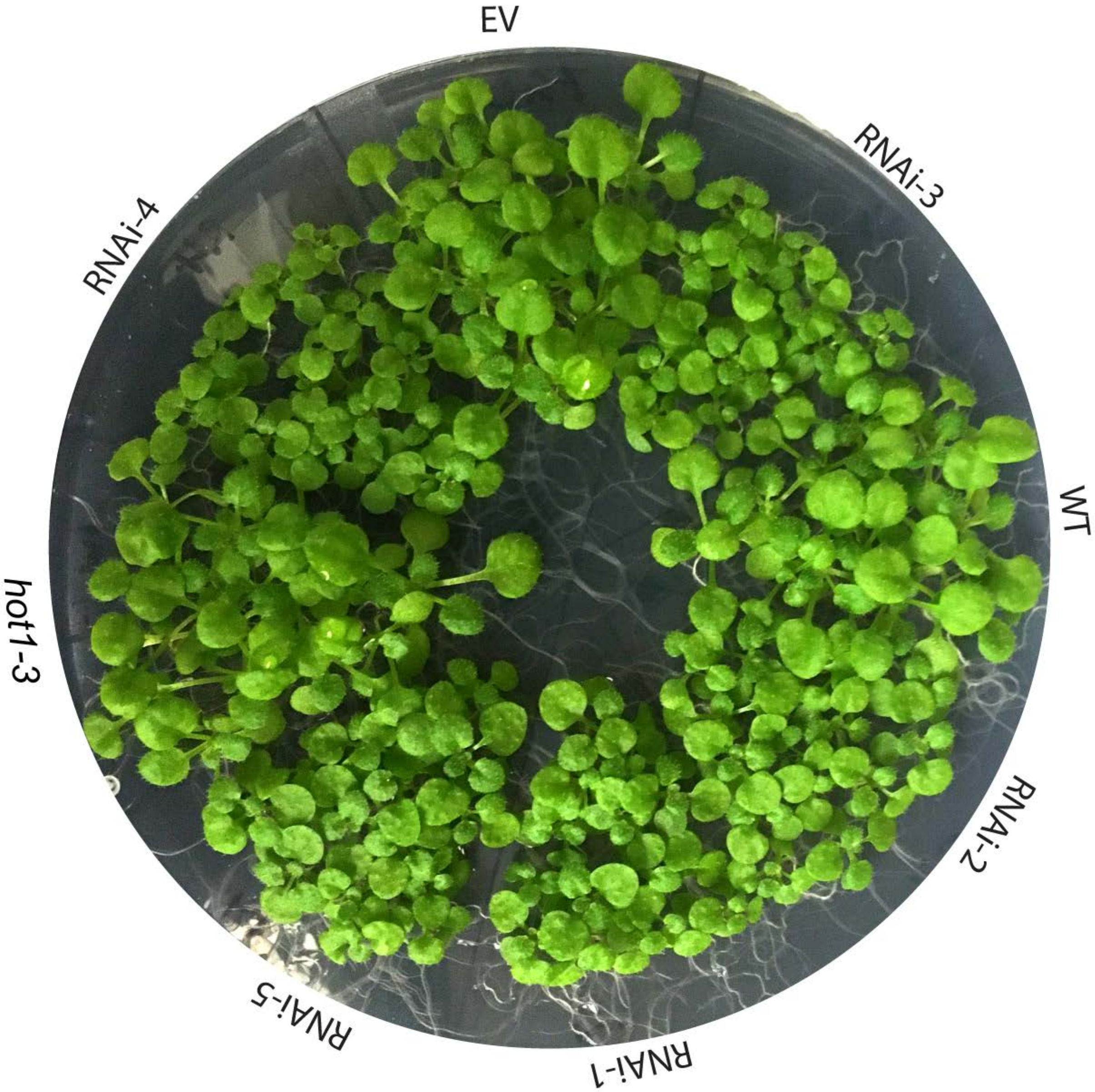


Figure S4. Seedlings of wild type (WT), empty vector control (EV), *hot1-3* and five *ATPd* RNAi lines grown on half-strength MS agar media for 17 days without heat stress.

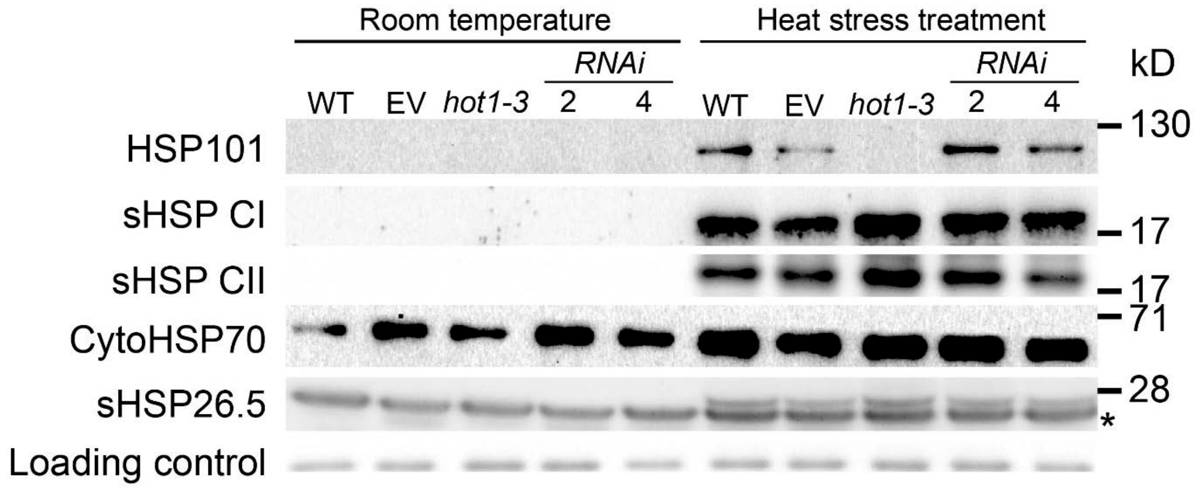


Figure S5. Protein levels of major cytosolic heat shock proteins (HSP) and a mitochondrial sHSP are not differentially regulated in *ATPd* RNAi lines. Total proteins were extracted from 10-day old seedlings of wild type (WT), empty vector control (EV), *hot1-3* and two RNAi plants treated with or without heat stress (38℃ for 1.5 hours followed by 2 h recovery at room temperature). All samples were probed with antibodies against cytosolic HSP101, HSP70, HSP70, sHSP class I, sHSP class II and mitochondrial sHSP26.5. An asterisk indicates non-specific bands. Non-specific bands from sHSP class II immunoblot were shown as a loading control. The experiments were repeated three times with similar results.

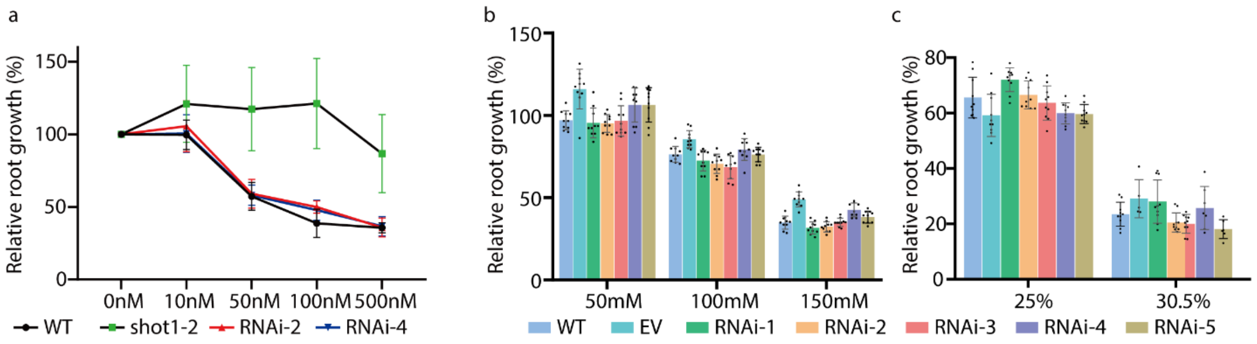


Figure S6. No significant differences in paraquat, salt and drought stresses were observed between *ATPd* RNAi lines and wild type (WT). Seedlings were grown on regular half-strength MS agar media for 4 days before transferring to media with or without chemicals. (a) Root growth for 3 days after transferring to paraquat containing plates was measured and shown as a relative growth to control plates. (b) Root growth for 6 days after transferring to NaCl containing plates was measured and shown as a relative growth to control plates. (c) Root growth for 6 days after transferring to PEG8000 containing plates was measured and shown as a relative growth to control plates. The paraquat, salt, and drought stresses were repeated three times, once and twice, respectively. Similar results were obtained for drought and paraquat stress treatments.

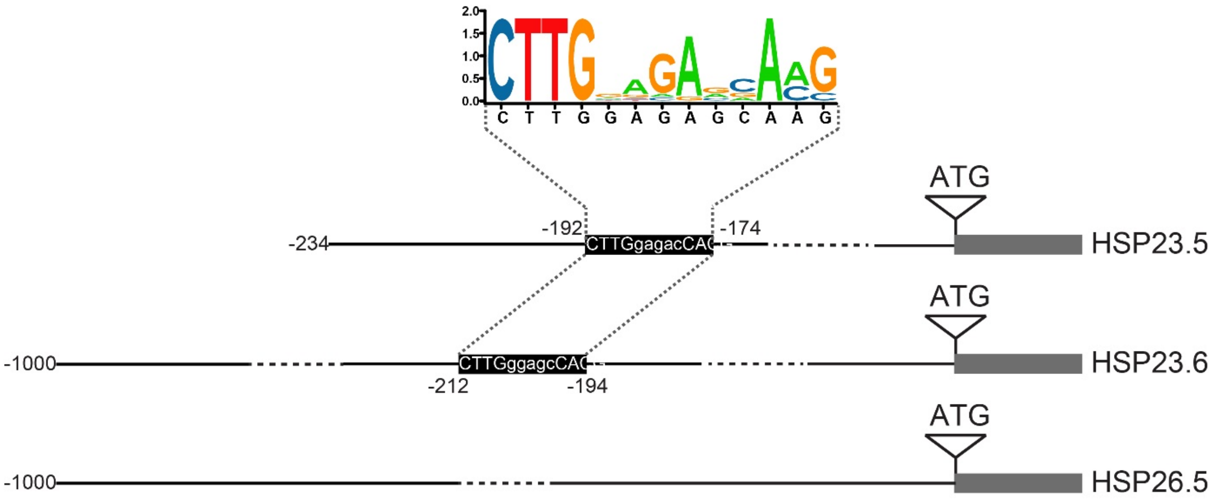


Figure S7. Mitochondrial dysfunction motif (MDM) exists in the promoter region of sHSP23.5 and sHSP23.6 while not in sHSP26.5. The MDM seqlogo was obtained from promoter sequences of AOX1a, UPOX1, MGE1, NDB4, OM66, HSP23.5, ANAC013, and SOT12 as previously identified (De Clercq et al., 2013). MDM consensus (CTTGNNNNNCA[AC]G) sequence was searched up to 1000 bp upstream of the start codons of HSP23.6 and HSP26.5.

**Supporting tables**

Table S1. Enrichment of different KEGG pathways from proteomics study comparing *ATPd* RNAi line with wild type.

| Type | Name | Size | Score | P value | Benj. Hoch. FDR | Mean | Median |
|---|---|---|---|---|---|---|---|
| KEGG pathway name | Glucosinolate biosynthesis | 7 | -0.6889 | 0.0016 | 0.0146 | -2.7162 | -2.5429 |
| KEGG pathway name | Sulfur metabolism | 14 | -0.5862 | 0.0002 | 0.0028 | -2.4974 | -1.6901 |
| KEGG pathway name | Starch and sucrose metabolism | 17 | -0.5065 | 0.0003 | 0.0043 | -1.8158 | -1.8536 |
| KEGG pathway name | Phagosome | 20 | -0.4274 | 0.0010 | 0.0098 | -1.3179 | -1.3681 |
| KEGG pathway name | 2-Oxocarboxylic acid metabolism | 35 | -0.3501 | 0.0004 | 0.0046 | -1.2374 | -1.6259 |
| KEGG pathway name | Biosynthesis of amino acids | 95 | -0.3337 | 0.0000 | 0.0000 | -1.1355 | -1.3257 |
| KEGG pathway name | Citrate cycle (TCA cycle) | 44 | -0.3095 | 0.0005 | 0.0049 | -1.0088 | -1.3266 |
| KEGG pathway name | Glycolysis / Gluconeogenesis | 52 | -0.3042 | 0.0002 | 0.0028 | -0.9686 | -1.3241 |

Table S2. Primers used in this study.

| *Gene identifier* | *Primer name* | *Primer sequence (5'->3')* | *Purpose* |
|---|---|---|---|
| *AT3G52300* | *ATPd-F1* | *GCCCACTACATTTCCAATTCC* | *Genotyping for atpd mutants* |
| *AT3G52300* | *ATPd-R1* | *GAGAGTTTTCTTTCCAAGCCG* | *Genotyping for atpd mutants, RT-PCR* |
| *AT3G52300* | *ATPd-R2* | *CGCAAGGCTCCATCAAGTAA* | *Genotyping for atpd-1 in complementation lines* |
| *AT3G52300* | *ATPd-F2* | *ATGAGCGGAGCCGGTAAGA* | *RT-PCR* |
| *AT3G52300* | *ATPd-F3* | *AATCTTGCTTAAACAGAGAATGGAT* | *Genomic DNA construct for complementation* |
| *AT3G52300* | *ATPd-R3* | *AGCATTGTGTCTTGAAGAACGA* | *Genomic DNA construct for complementation* |
| *AT3G52300* | *ATPd-F4* | *caccAAAGCTCAGCACCATGACTG* | *RNAi, pMK118 plasmid* |
| *AT3G52300* | *ATPd-R4* | *TCACTGAAGAACGAAACGAAACA* | *RNAi, pMK118 plasmid* |
| *AT3G52300* | *ATPd-F5* | *gactctagaggcgcgccAAAGCTCAGCACCATGACTG* | *RNAi, pMK124 and pMK127 plasmids* |
| *AT3G52300* | *ATPd-R5* | *gacggatccatttaaatTCACTGAAGAACGAAACGAAACA* | *RNAi, pMK124 and pMK127 plasmids* |
| *AT4G05320* | *UBQ10-F* | *GATCTTTGCCGGAAAACAATTGGAGGATGGT* | *RT-PCR* |
| *AT4G05320* | *UBQ10-R* | *CGACTTGTCATTAGAAAGAAAGAGATAACAGG* | *RT-PCR* |
| *NA* | *SAIL-LB3* | *TAGCATCTGAATTTCATAACCAATCTCGATACAC* | *T-DNA specific primer for genotyping of atpd mutants* |
| *AT3G18780* | *ACT2_F* | *ATCGAGAAGAACTATGAATTAC* | *qRT-PCR* |
| *AT3G18780* | *ACT2_R* | *AAGTGCTGTGATTTCTTT* | *qRT-PCR* |
| *AT3G22370* | *AOX1a_F* | *AGCATCATGTTCCAACGACGTTTC* | *qRT-PCR* |
| *AT3G22370* | *AOX1a_R* | *GCTCGACATCCATATCTCCTCTGG* | *qRT-PCR* |
| *AT3G22360* | *AOX1b_F* | *CCGTGAAATCTCTTCGATGGC* | *qRT-PCR* |
| *AT3G22360* | *AOX1b_R* | *CATCCCTCCAACCATTCCTG* | *qRT-PCR* |
| *AT3G27620* | *AOX1c_F* | *GTTCCTTCAACTTTACCGGAC* | *qRT-PCR* |
| *AT3G27620* | *AOX1c_R* | *CGTCTCTAGCATAATCGCTC* | *qRT-PCR* |
| *AT2G21640* | *UPOX1_F* | *CAAACCTCAAGGATCACATGGATGA* | *qRT-PCR* |

| | | | |
|---|---|---|---|
| *AT2G21640* | *UPOX1_R* | *GCCTTGGAGAAGCTCCCGAATATCT* | *qRT-PCR* |
| *AT1G32870* | *ANAC013_F* | *CCATAGAGGCAGGGCACCTAATGG* | *qRT-PCR* |
| *AT1G32870* | *ANAC013_R* | *CCATTCTTAGGACCAGACCCAC* | *qRT-PCR* |
| *AT2G03760* | *SOT12_F* | *ATCGCATCTTTCGCTCCC* | *qRT-PCR* |
| *AT2G03760* | *SOT12_R* | *TCCGCTTCATCTCAACTTCG* | *qRT-PCR* |
| *AT5G55200* | *MGE1_F* | *ACGCAGTGTTCCAAGTCCCA* | *qRT-PCR* |
| *AT5G55200* | *MGE1_R* | *TCTTTGCCGCCTTCTTGATT* | *qRT-PCR* |
| *AT2G20800* | *NDB4_F* | *TCAAGTCTTAAGGGCACAACA* | *qRT-PCR* |
| *AT2G20800* | *NDB4_R* | *CGGAAGAGAGAGGCTCTCTCG* | *qRT-PCR* |
| *AT3G50930* | *OM66_F* | *TGCTGAGACCAGGACGTATG* | *qRT-PCR* |
| *AT3G50930* | *OM66_R* | *ACCTTCCTCGATCTTGCTGA* | *qRT-PCR* |
| *AT3G11630* | *2CPA_F* | *CCCAACAGAGATTACTGCCT* | *qRT-PCR* |
| *AT3G11630* | *2CPA_R* | *ATAGTTCAGATCACCAAGCCC* | *qRT-PCR* |
| *AT5G08670, AT5G08680, AT5G08690* | *ATP-β_F* | *CGTGCCCGTAAGATCCAG* | *qRT-PCR* |
| *AT5G08670, AT5G08680, AT5G08690* | *ATP-β_R* | *TCGATACCTCCAACCATG* | *qRT-PCR* |
| *AT5G47030* | *ATP-δ_F* | *GGGATCCTCATTTGCGGTGTCATTC* | *qRT-PCR* |
| *AT5G47030* | *ATP-δ_R* | *AACTGCAGGGCGGCGATTTGG* | *qRT-PCR* |
| *AT3G52300* | *ATPQ-q1* | *TTGATGCTTTGTTGGTGGAAC* | *qRT-PCR* |
| *AT3G52300* | *ATPQ-q2* | *ACCGTTCAGACTCCTTGAGC* | *qRT-PCR* |
| *AT5G51440* | *23.5_F* | *TCAAACCGACATGTTTCTCG* | *qRT-PCR* |
| *AT5G51440* | *23.5_R* | *AAGCTTCTCGTTGGAGTAAACG* | *qRT-PCR* |
| *AT4G25200* | *23.6_F* | *GTCGTCGTAGCTCCTAACGG* | *qRT-PCR* |

| | | | |
|---|---|---|---|
| *AT4G25200* | *23.6_R* | *CAAATGGCATCTGCTCTCGC* | *qRT-PCR* |
